# Pathogen-Induced Damage in Drosophila: Uncoupling Disease Tolerance from Resistance

**DOI:** 10.1101/2025.01.31.635923

**Authors:** Priscilla A. Akyaw, Tânia F. Paulo, Elvira Lafuente, Élio Sucena

## Abstract

Immune response against infections can be divided into mechanisms of resistance that ensure active pathogen elimination, and mechanisms of disease tolerance, which include processes that return the host to physiological homeostasis without involving direct pathogen control. Studies on host immune responses to infection have mostly targeted mechanisms of resistance, and consequently, these are now well-described in both vertebrates and invertebrates. By comparison, the mechanistic basis of disease tolerance is less well understood. This is in part because both processes interact and can be difficult to separate under an infection scenario. Using the highly tractable insect model *Drosophila melanogaster* exposed to its natural entomopathogen, *Pseudomonas entomophila*, we aimed to tease apart mechanisms of disease tolerance from those of resistance. To this aim, we reasoned that oral exposure to heat-killed entomopathogenic bacteria should require disease tolerance without relying on resistance. Using this method, we observe that oral exposure to heat-killed *P. entomophila* causes mortality and reduced fecundity in *D. melanogaster*. We confirm that this reduction in fitness-related traits depends on the duration of the exposure, is sexually dimorphic, and is dependent on the virulence of the bacterium. We also found the microbiota to play a role, with its presence exacerbating the deleterious effect on host survival. This experimental framework, which may be extended to other systems, can be instrumental towards an understanding of the molecular, genetic, and physiological basis of disease tolerance and its interactions with resistance mechanisms.

## Introduction

The ability to overcome an infection without enduring deleterious fitness consequences strongly influences the evolution of both hosts and pathogens (Hite et al., 2023; Sadd & Schmid-Hempel, 2009). Hosts employ a variety of countermeasures against pathogens including behavioural avoidance (e.g. staying clear of contaminated food, water, and other infected hosts) (Hart & Hart, 2018; Lopes et al., 2022; Sarabian et al., 2018; Zélé et al., 2019), and the classical immunological responses involving mechanisms of resistance (i.e. the use of immune effectors to eliminate or reduce pathogen load) and of disease tolerance (i.e. the minimizing of the effects of damage caused by the infection process) (Ayres, 2017; Råberg et al., 2009; Schmid-Hempel, 2003; Schulenburg et al., 2008). These strategies ultimately ensure the prevention and control of infection to promote host survival and reproduction (Kutzer & Armitage, 2016; Mccarville & Ayres, 2017; Råberg et al., 2007).

In the last two decades, *D. melanogaster* has been instrumental in further characterizing these two arms of the immune response (Aggarwal & Silverman, 2008; Ayres et al., 2008; Chapman et al., 2020; De Gregorio et al., 2002; Howick & Lazzaro, 2017; Lemaitre & Hoffmann, 2007; Paulo et al., 2023; Valanne et al., 2011; Westlake et al., 2024). A pivotal moment was the discovery of the involvement of the Toll and IMD pathways in the mechanisms of resistance that target invading pathogens for elimination, through the regulation of antimicrobial peptides (AMPs) and reactive oxygen species (ROS) production (De Gregorio et al., 2002; Lemaitre et al., 1997; Sun & Faye, 1992). These and subsequent studies have uncovered the functional roles of specific AMPs such as Cecropins (*Cec*) (Hultmark et al., 1983), Diptericin (*Dpt*) (Lambert et al., 1989), Defensin (*Def*) (Cociancich et al., 1993) and Drosomycin (*Drs*) (Fehlbaum et al., 1994), as well as the dynamics of their transcriptional activation during infection. More recently, research has enquired on how these processes lead to pathogen clearance (Lemaitre et al., 1997; Schlamp et al., 2021) and/or pathogen containment at levels that prevent harm (Acuña Hidalgo et al., 2022; Duneau & Ferdy, 2022; Troha & Buchon, 2019), as well as on the regulatory mechanisms that govern them (Aggarwal & Silverman, 2008; Chambers & Schneider, 2012; Lazzaro & Tate, 2022; Vincent & Dionne, 2021).

In contrast, disease tolerance still awaits a comparably deep mechanistic characterization. It is expected that the processes involved include the maintenance of homeostatic conditions and normal physiological functions by repair and/or prevention of damage caused during infection (Ayres & Schneider, 2012; Baucom & de Roode, 2011; Martins et al., 2019; Medzhitov et al., 2012; Råberg, 2014). Operationally, disease tolerance is measured as the relative status of a health criterion upon infection or of a proxy measure of immune elicitors/effectors (Ayres & Schneider, 2012; Best et al., 2008; Kutzer et al., 2023; Prakash et al., 2022; Råberg et al., 2009; Råberg et al., 2007; Råberg, 2014; Simms, 2000). However, even the seemingly simple choice of what criterion should be considered and what parameters measured to assess physiological state, is highly variable and encompasses distinct phenotypes and terminology that include “fitness” (Kutzer & Armitage, 2016; Medzhitov et al., 2012; Simms, 2000), “health”, and “performance” (Doeschl-Wilson et al., 2012; Råberg et al., 2009; Råberg et al., 2007; Regoes et al., 2014). Moreover, resistance and disease tolerance are likely to be mechanistically and evolutionarily intertwined (Ayres, 2020b; Ayres & Schneider, 2008; Best et al., 2008, 2014; Duneau et al., 2024; Kutzer et al., 2018; Kutzer & Armitage, 2016; Paulo et al., 2023; Prakash et al., 2023; Råberg et al., 2009; Råberg et al., 2007). Ultimately, the knowledge gap regarding the mechanisms of disease tolerance is partially explained by the difficulty in disentangling them from resistance in a solid operational way, allowing for focused experimental approaches (Ayres, 2020b; Duneau et al., 2017; Råberg et al., 2009; Råberg et al., 2007; Simms, 2000).

One first step in unravelling the mechanisms underlying disease tolerance has been to distinguish mortality tolerance, defined as higher survival under comparable pathogen loads, from fecundity tolerance, understood as maintaining reproductive output during infection (Roy & Kirchner, 2000; Vale & Little, 2012; Kutzer & Armitage, 2016). Studies focusing on mortality tolerance have uncovered a number of genes, including *Eiger*, *CrebA*, *Pirk*, and *IRC*, involved in promoting host survival upon systemic infection without affecting pathogen load (Brandt et al., 2004; Kleino et al., 2008; Lissner & Schneider, 2018; Troha et al., 2018). Yet, the specific ways in which these genes shape this response and the nature of the physiological processes in which they participate, are not fully understood.

To gain a deeper understanding of disease tolerance, it is fundamental to characterize the consequences of infection on other important life-history traits, such as reproduction, stress response, growth, tissue repair capacity, as well as energy acquisition and allocation (Arbuthnott, 2018; Ayres, 2020a; Best et al., 2014; Bonneaud et al., 2019; Kotas & Medzhitov, 2015; Roy & Kirchner, 2000; Stearns & Stearns, 1992). Furthermore, it is also critical to understand whether such processes and mechanisms promoting disease tolerance are general or deployed differently, according to variations in challenge such as distinct pathogens or routes of infection (Gupta, Vasanthakrishnan, et al., 2017; Martins et al., 2013). Be as it may, disease tolerance also encompasses *sensu latu* response to damages, whether directly inflicted by the pathogen or self-inflected (Ayres & Schneider, 2012; Brandt et al., 2004; Lazzaro & Tate, 2022; Medzhitov et al., 2012; Schneider & Ayres, 2008; Soares et al., 2014).

In insects, there is evidence suggesting that immunopathology contributes to mortality during infection (Brandt et al., 2004; Lazzaro & Tate, 2022; Li et al., 2020; Prakash et al., 2024; Pursall & Rolff, 2011; Sadd & Siva-Jothy, 2006). In particular, previous work in *D. melanogaster* established that immune activation is achieved by the recognition of microbial cell wall components and also by virulence factors secreted by specific pathogens (Brandt et al., 2004; Chamy et al., 2008; Dieppois et al., 2015; Opota et al., 2011). Therefore, *D. melanogaster* is likely to sustain these two non-mutually exclusive types of fitness costs when it is infected: indirect damage induced by immunopathology and direct damage imposed by the pathogen (via secretion of virulence factors and toxins). Responding to either or both types of damage will require mechanisms of disease tolerance.

To understand how disease tolerance promotes host fitness independently from the action of resistance in reducing pathogen load, we measured survival and fecundity of *D. melanogaster* after exposure to an inactivated form of *Pseudomonas entomophila*. This gram-negative bacterium is a natural entomopathogen of *D. melanogaster* that, ensuing the detection of virulence factors and/or microbial cell wall components, activates both local and systemic immune responses at different life stages (Dieppois et al., 2015; Lemaitre, 2015). Although *Drosophila* deploys a very strong immune response upon oral infection, mainly through the upregulation of specific AMPs and ROS towards this bacterium (Ha et. al. 2005), it suffers severe damage to the gut epithelium leading to high mortality within the first 24 hours after exposure (Liehl et al., 2006; Troha et al., 2018; Vodovar et al., 2005).

In this work, we measured the mortality and fecundity tolerance exhibited by *D. melanogaster* when fed with heat-killed *P. entomophila*. We reasoned that the host response to oral exposure to heat-killed pathogenic bacteria should require disease tolerance without relying on resistance since there is no pathogen proliferation to control. We first characterized how feeding on heat-killed bacteria affected host survival and fecundity, and quantified damage to the gut epithelium after this treatment. Additionally, we measured gene expression profiles of key components of the *Drosophila* immune response, such as canonical AMPs, and components of the ROS and stress response pathways. In this way, we have established a new experimental framework with which to measure disease tolerance to pathogen-derived damage that allows for the disentangling of these effects from their immune-resistance counterparts. With this approach, we hope to contribute to coming one step closer towards the mechanistic basis for disease tolerance in an oral infection context.

## Results

### 1. Establishing a protocol to measure disease tolerance

The gram-negative bacterium *Pseudomonas entomophila* is highly virulent to *Drosophila melanogaster*, killing more than 50% of flies within 48h after ingestion (Lemaitre, 2015; Vallet-Gely et al., 2010). To determine whether oral exposure to food containing heat-killed (HK) *P. entomophila,* would induce a measurable fitness cost in *D. melanogaster,* we fed it to adults for periods of two, three, four, or five days (Fig. S1A), after which, we measured survival (Fig. 1A) and fecundity (Fig. 1B) for 12 days. We set the heat inactivation treatment to 55°C as this was the lowest temperature for which we did not observe bacterial colonies upon subsequent plating.

**Figure 1.**
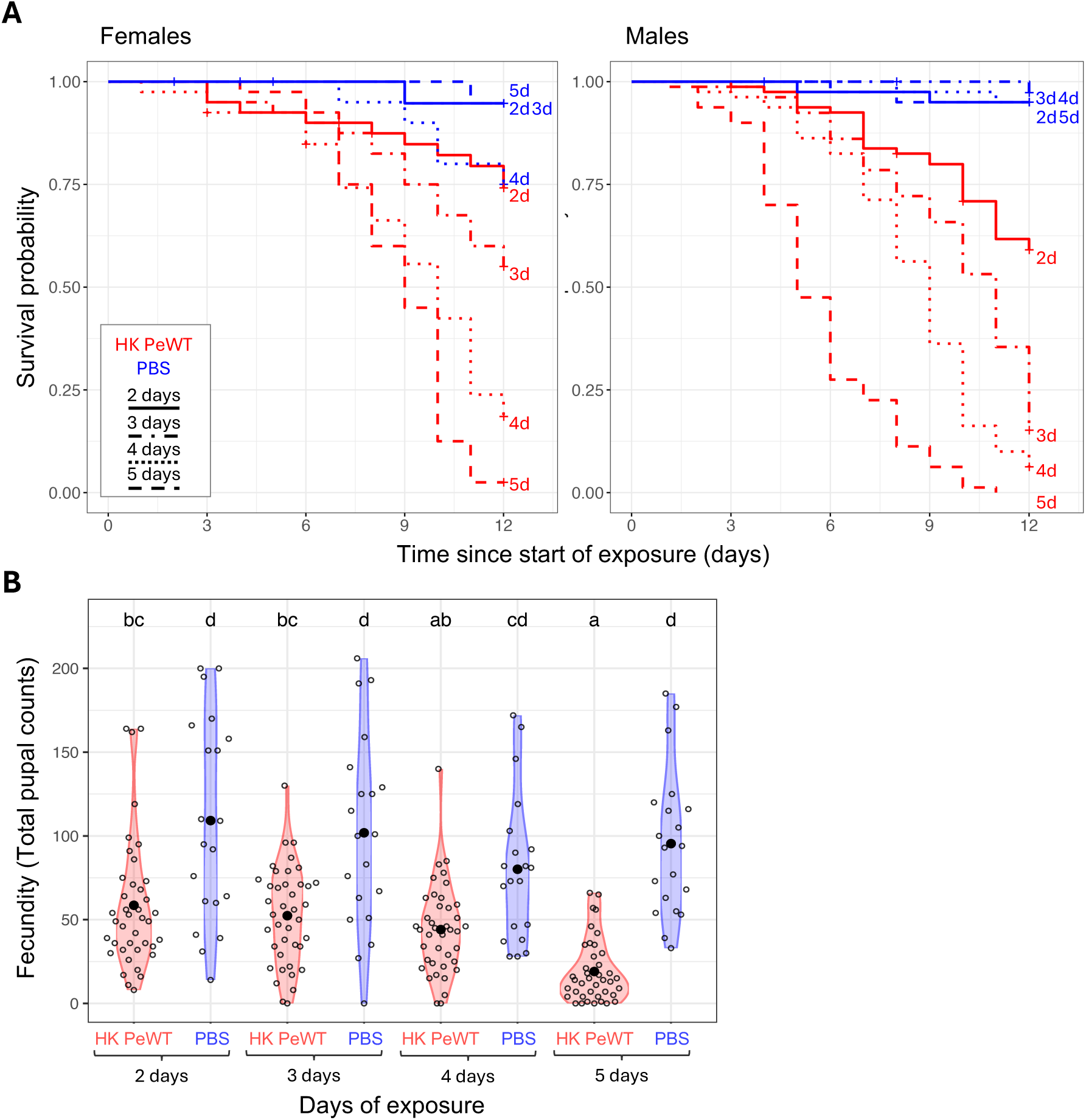
Exposure to heat-killed *P. entomophila* entails survival and fecundity costs. Survival and fecundity were monitored for 12 days in flies exposed to heat-killed (HK) *P. entomophila* (HK PeWT - red) or control food (PBS - blue). **A)** Survival curves for females (left panel) and males (right panel) exposed for two, three, four, or five days. HK *P. entomophila* treatment led to increased mortality as compared to PBS, independently of the number of days of exposure (p<0.05 for all comparisons), except for females in the two-day exposure group. **B)** Female reproductive output measured as total pupal counts. Flies exposed to HK *P. entomophila* have consistently lower reproductive output compared to control, with the 5-day exposure regime showing a significantly stronger effect than the other exposure times. Differences between groups were estimated by post hoc comparisons (Tukey’s honest significant differences) and are indicated by different letters in each plot (p-value<0.05).

We observed a significant effect of treatment on mortality that was dependent on sex (Anova(Cox): χ²(1)= 8.9, p= 0.003) and on the length of exposure (Anova(Cox): χ²(3)= 10.13, p= 0.02) (Fig. 1A, Tables S1, S2). Except for females from the two-day exposure treatment (EMMs(Cox): est= -1.74 (SE= 1.05), p= 0.097), all flies exposed to food mixed with HK *P. entomophila* were more susceptible than those exposed to food mixed with PBS (p< 0.05 in all cases), and the longer the exposure, the highest the mortality (Fig. 1A) (Table S2). Flies exposed for two days had the highest survival rates, with 74% for females and 59% for males. The three-day exposure group showed a steeper survival decline: females at 55% survival and males at only 15%. The five-day exposure resulted in the lowest survival, with only 2% for females and no surviving males by day 12. There was a significant difference between the number of exposure days for all paired combinations (p< 0.01 in all cases) except for females exposed for two and three days (EMMs(Cox): est= -0.69 (SE=0.39), p= 0.3) and between four and five days (EMMs(Cox): est= -0.54 (SE= 0.25), p= 0.13) (Fig. 1A, Table S2).

These results reveal also a strong sexual dimorphism in the response to oral exposure to HK *P. entomophila*, with males having a significantly higher risk of death (approximately twice) compared to females (Cox: HR= 2.12, p< 0.001).

With regards to fecundity, we also found it to be significantly reduced in flies exposed to HK *P. entomophila* as compared to the control group (Anova(lmer): χ²(1)= 23.21, p= 1.45e-06) (Fig. 1B), with a tendency to correlate negatively with the length of exposure, though only significant when comparing four and five days to two and three days exposures (Fig. 1B, Table S2). The effect of exposure to HK *P. entomophila* led to an average reduction in fecundity of about 55% relative to PBS control (i.e. mean fecundity estimates were 96.6 and 43.5 for the HK and the PBS treatments, respectively) (Fig. 1B, Table 5).

Having shown that feeding heat-killed *P. entomophila* negatively impacted both survival and fecundity in adult flies, we sought to generalize this effect by testing its independence from the bacteria inactivation method itself. To that aim, we first inactivated the bacteria with an alternative method consisting of incubation in paraformaldehyde (PFA) (Fig. S2A). PFA inactivation affected host survival negatively (Table S1) compared to the control group (females: EMMs(Cox): est= -2.40 (SE= 0.27), p= 5.33e-14), showing a similar tendency to that of the heat-killing protocol for both sexes (Fig. S2A, Table S3). Secondly, to gain insight into the relationship between the observed effects on survival and the level of denaturation of the bacteria, we tested a harsher heat inactivation temperature of 95°C. Analyses of survival upon exposure to the two heat-killing temperatures, revealed a significant effect of this variable (Table S1) most notably in females (i.e. significant temperature by sex interaction) (Anova(Cox): χ²(2)= 26.8, p=1.54e-06), where inactivation at 95 °C led to considerably more mortality than inactivation at 55 °C (Fig. S2B, Tables S1, S3).

In conclusion, exposure to inactivated *P. entomophila* led to increased mortality and reduced female fecundity (shown for heat-inactivated bacteria), both scaling positively with the duration of exposure. Based on these findings, all subsequent experiments were performed under a protocol using a three-day exposure to 55°C heat-killed *P. entomophila*.

### 2. Mortality after exposure to heat-killed bacteria depends on its pathogenicity

To determine whether the fitness costs observed in flies after exposure to heat-killed bacteria were specific to the entomopathogen *P. entomophila* or if they reflected a general response, we tested additional bacterial species through the same protocol. For that, we followed survival of flies fed with four different HK gram-negative bacteria with described varying levels of pathogenicity through the oral route. We used *Pseudomonas putida*, an avirulent bacterium closely related to *P. entomophila* (Vodovar et al., 2006), *Pectobacterium carotovorum* (formerly known as *Erwinia carotovora carotovora*) (strain Ecc-15) which is known to secrete virulence factors but does not induce high mortality in *Drosophila* (Basset, 2000; Buchon, Broderick, Poidevin, et al., 2009), a mutant version of this bacterium in which the virulence factors have been deleted (strain Ecc-71), and *Escherichia coli (E-coli)* K-12 strain which is considered non-pathogenic to *D. melanogaster* (Nixon et al., 2024; Raval et al., 2023).

Host survival was not significantly affected upon exposure to these heat-killed bacterial species for three days except for *P. entomophila*, which led to mortality exceeding that of the PBS control (p>0.05 for all comparisons) (Fig. 2A, Tables S1, S4) with a sex-dependent effect (Anova(Cox): χ²(5)= 16.01, p= 0.007).

**Figure 2.**
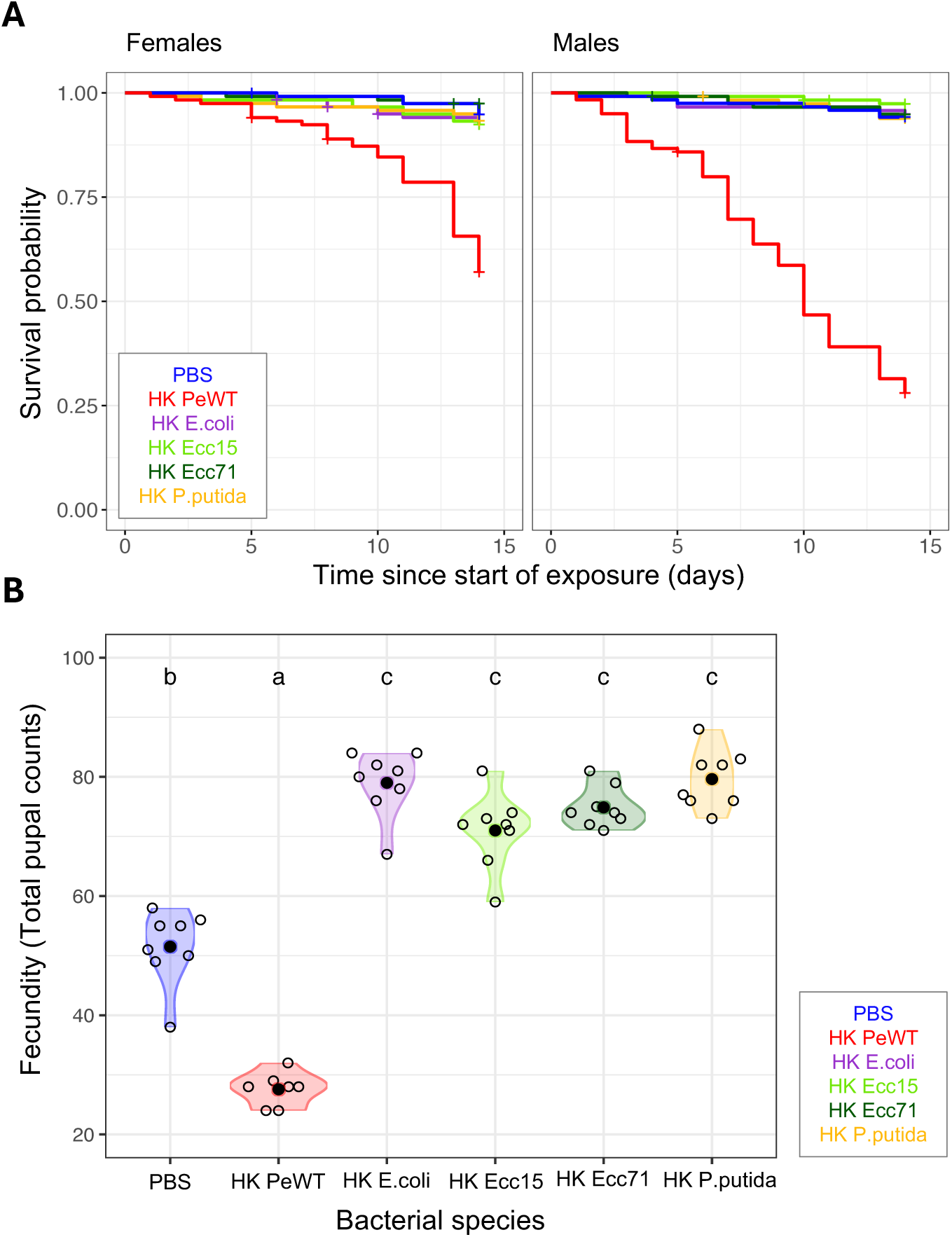
Survival and fecundity upon exposure to heat-killed (HK) bacteria depend on bacterial virulence. Survival and fecundity of flies exposed to PBS control food (PBS – blue) or to heat-killed (HK) bacteria with different levels of virulence: *P. entomophila* (PeWT - red), *Escherichia coli* (E.coli - purple), *Erwinia carotovora carotovora* 15 (Ecc15 - light green), *E. carotovora carotovora* 71 (Ecc71 – dark green) or *Pseudomonas putida* (P.putida - yellow). **A)** Survival curves in females (left panel) and males (right panel) over 14 days shows that there is no difference in survival between the PBS-exposed group and all other bacteria exposed flies, except for *P. entomophila* (p<0.05 in all cases) (Tables S1, S4). **B)** Mortality-corrected reproductive output measured as cumulative daily pupal count over 14 days (see Material and Methods). Reproductive output is lowest upon exposure to heat-killed *P. entomophila*, followed by PBS. Exposure to all other bacteria species leads to increased reproductive output (p<0.05 in all cases). Differences between groups were estimated by post hoc comparisons (Tukey’s honest significant differences) and are indicated by different letters in each plot (p-value<0.05).

Concerning female fecundity, we found it to be significantly affected by exposure to the different bacterial species (Anova(glmm): χ²(5)= 256.34, p<2.2e-16, Table S1) and different from the PBS control (p<0.05 in all cases, Table S4). Notably, all females exposed to non-virulent bacteria showed a higher fecundity, equivalent amongst them, than those fed with PBS (p< 0.05 for all pairwise comparisons, Table S4). As expected, and previously shown, females exposed to the HK *P. entomophila* displayed a significant decrease in fecundity as compared to control (EMMs(glmm): est= -0.66 (SE= 0.08), p= 1.45e-13) (Fig. 2B, Table S4).

### 3. Virulence factors of heat-killed *P. entomophila* are necessary to induce mortality and fecundity costs

To confirm that the virulence factors of *P. entomophila* cause fitness costs under the HK exposure treatment, we measured survival and fecundity after feeding flies with either wild-type bacteria (PeWT) or the mutant avirulent strain *P. entomophila* ΔGacA, which carries a Tn5 mini transposon in the GacA gene (hereafter, PeGacA) (Vodovar et al., 2005).

In contrast to exposure to HK PeWT, survival was higher in flies exposed to HK PeGacA treatment in both males and females (Fig. 3A, Table S1). The PeGacA strain of the bacterium failed to induce higher mortality than the PBS control in either sex (EMMs(Cox) with p>0.05, Table S5) (Fig. 3A). Once more, we observed strong effects of exposure to HK PeWT on survival, whereby both sexes die more than in the PBS control (Fig. 3A, Table S1, S5), more prominently in males (EMMs(Cox): est= 4.18 (SE= 1.1), p= 0,0004) than in females (EMMs(Cox): est= 3.2 (SE= 1.1), p= 0.01), (Table S5). We extended this experiment to 17 days to evaluate potential delayed effects; however, the survival curves showed relatively steady slopes over the following second and third weeks, and therefore, subsequent experiments were conducted for 14 days.

**Figure 3.**
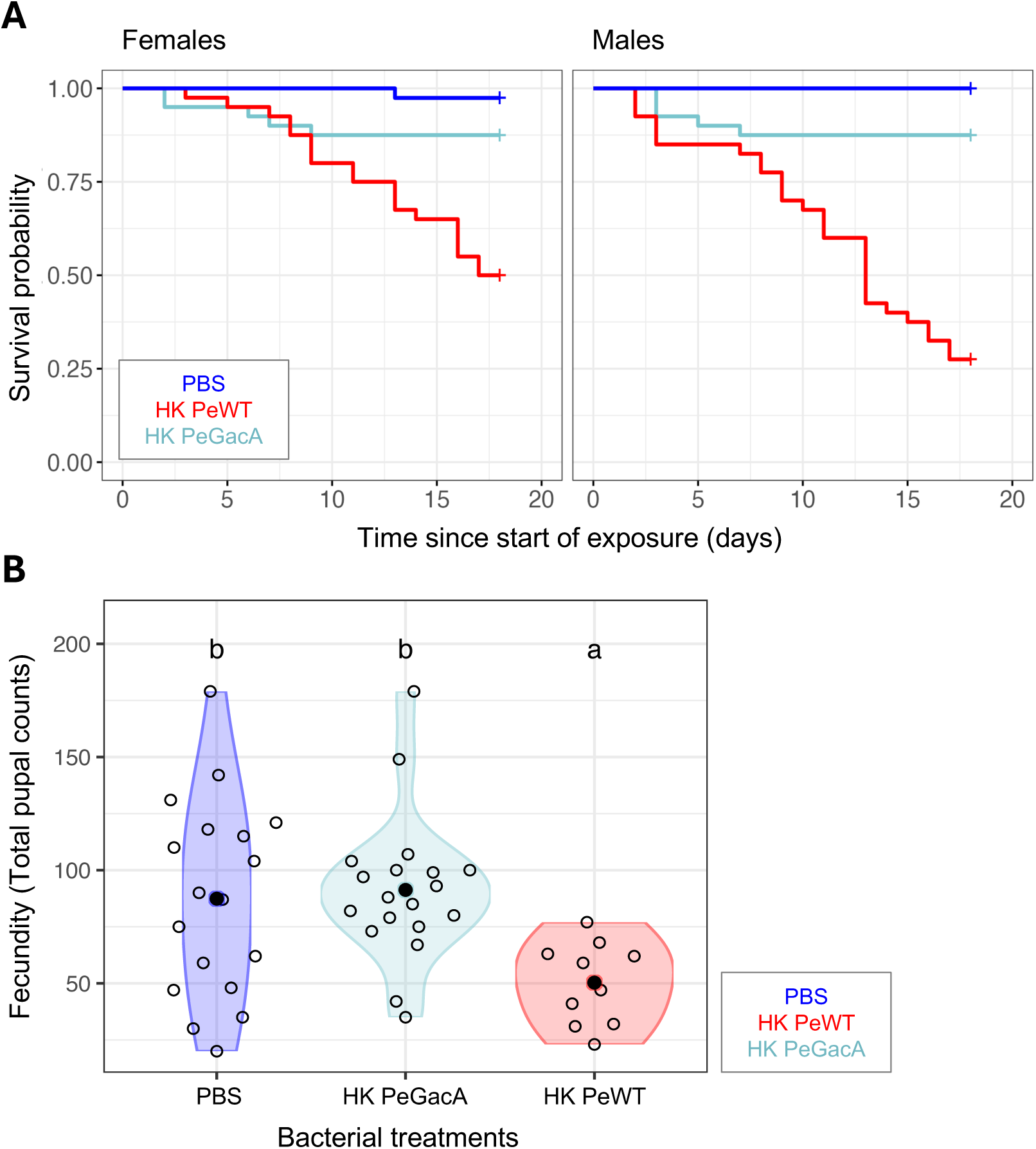
Exposure to a heat-killed non-virulent strain of *P. entomophila* does not affect mortality nor fecundity. Survival and fecundity in flies exposed to food containing heat-killed wild-type *P. entomophila* (PeWT - red), the avirulent *P. entomophila ΔGacA* mutant (PeGacA - light blue), or PBS for control (PBS - dark blue). **A)** Survival curves of females (left panel) and males (right panel) over 18 days show no difference in survival between the PBS- and GacA-exposed groups (p>0.05 in both sexes), in contrast with WT-exposed flies which are significantly different for all comparisons (p<0.01). As previously shown, response to WT *P. entomophila* differed between sexes (p<0.01). **B)** Mortality-corrected reproductive output measured as cumulative daily pupal count (see Material and Methods) shows that reproductive output in HK *P. entomophila* GAC- and PBS-exposed flies do not differ, and are both higher than in HK *P. entomophila* WT exposed flies. Differences between groups were estimated by post hoc comparisons (Tukey’s honest significant differences) and are indicated by different letters in each plot (p<0.05).

We also observed differences in fecundity between bacterial mutants (Anova(glmm): χ²(2)= 32.65, p= 8.15e-08). Flies exposed to HK PeGacA did not show a reduction in fecundity as the one observed with HK PeWT (Fig. 3B). In fact, fecundity in flies exposed to HK PeGacA was similar to that of flies exposed to PBS control and higher than under exposure to HK PeWT (Fig. 3B, Table S5).

In parallel, and as a somewhat positive control and confirmation of the pathogenesis of the bacteria, we ran the same experiments described above with live *P. entomophila* (Fig. S3, Table S1, S6). Live Infection with the wild-type *P. entomophila* led to mortality of over 90% within five to six days of infection (Fig. S3), as previously reported (Lemaitre, 2015; Neyen et al., 2014; Paulo et al., 2023; Vallet-Gely et al., 2010). We found a significant difference in survival between live PeWT, but not PeGacA, and sucrose treatments (Anova(Cox): χ²(2)= 162.6, p= 5.02e-36) and between sexes (Anova(Cox): χ²(1)= 4.47, p= 3.46e-02) (Fig. S3A, Tables S1, S6). Fecundity differed between treatments (Anova(glmm): χ²(2)= 157.18, p= 7.40E-35; Table S1), with the highest fecundity observed in the presence of live GacA, followed by the sucrose control, and the lowest fecundity in the presence of live PeWT (Fig. S3B, Tables S1, S6).

### 4. Exposure to heat-killed *P. entomophila* causes gut damage

The virulence factors of *P. entomophila* are known to induce severe gut epithelium damage during an oral infection (Lemaitre, 2015; Nonaka et al., 2020; Opota et al., 2011). Having found that exposure to heat-killed *P. entomophila* caused reduced survival and reproduction in a virulence factor-dependent manner (Figs 1, 2 and 3), we asked if this effect could be downstream of damage inflicted to the gut, as described for infections with live bacteria (Buchon, Broderick, Chakrabarti, et al., 2009; Liehl et al., 2006; Vodovar et al., 2005). To get a direct measure of cell damage, we looked at the number of apoptotic and mitotic cells using immunohistochemistry staining against genes Dcp-1 and PH3, respectively, in the posterior midgut region of individuals exposed to HK PeWT, HK PeGacA, or PBS (Ohlstein & Spradling, 2006; Zhai et al., 2018) (Fig. 4A).

**Figure 4.**
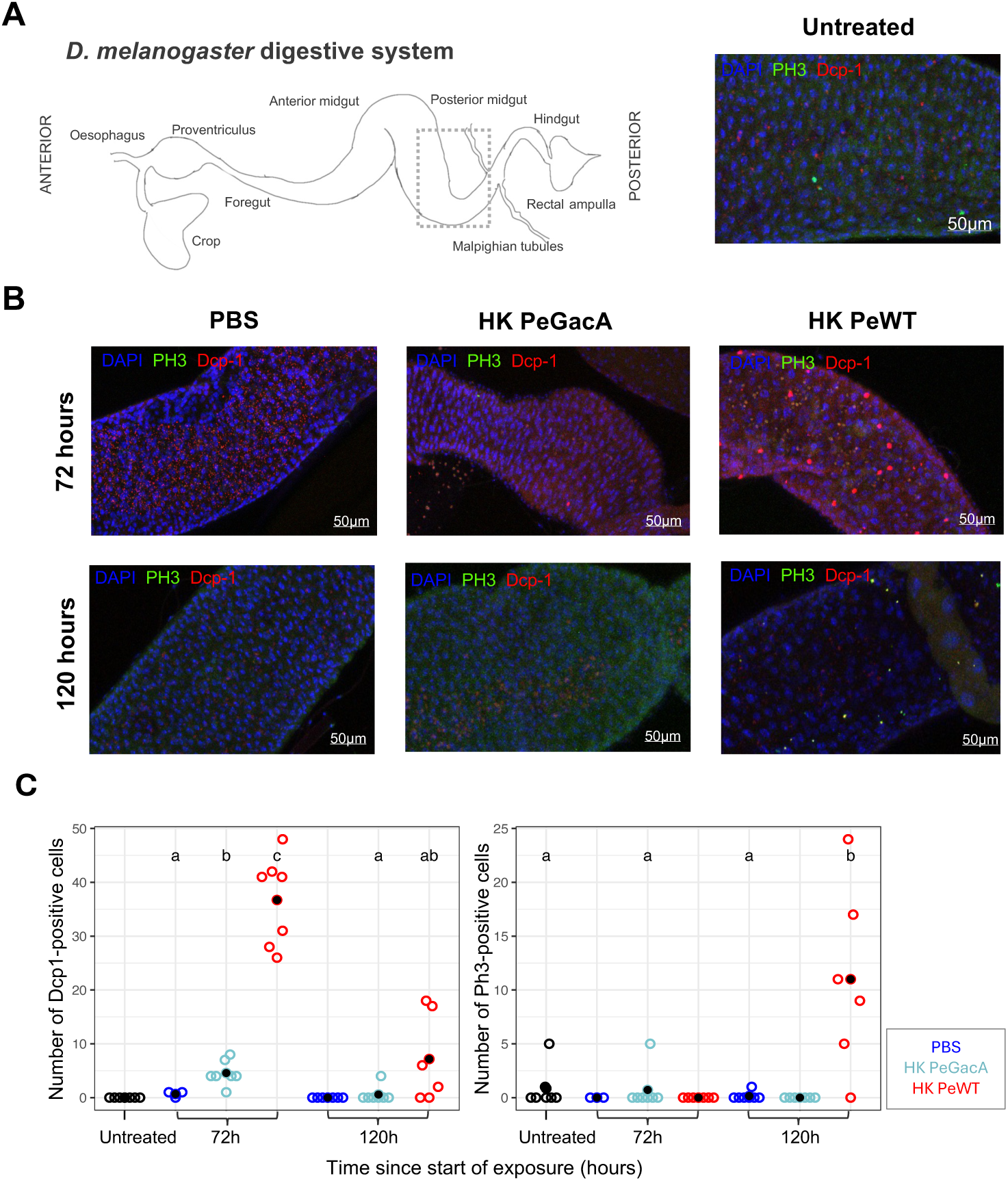
Feeding on heat-killed *P. entomophila* increases apoptosis and mitotic cell division in gut epithelial tissues. **A)** On the left, a cartoon depicting the *Drosophila* digestive system (anterior to the left). The dashed inset box delimits the midgut region shown in subsequent images of Immunofluorescent staining (A and B). Immunostaining shows antibodies against *Drosophila* caspase 1 (Dcp-1) in red and Phospho-H3 (PH3) in green, merged with DAPI in blue, as exemplified on the right panel by a picture from a control “untreated” sample. **B)** Guts of flies exposed to PBS, heat-killed ΔGacA mutant (HK PeGacA), or heat-killed wildtype (HK PeWT) *P. entomophila* at 72 hours (top) and 120 hours (bottom) after exposure. **C)** Quantification of Dcp-1 or PH3-positive cells per posterior midgut of males (N= 6 to 8). Exposure to HK PeGacA provoked a significant increase in apoptotic cells at 72 hours (p<0.05), which dissipated 2 days later. Feeding on HK PeWT led to a large increase in the number of Dcp-1-positive cells (p<0.01), which remained tendentially elevated at 120 hours. Mitotic gut cell numbers were significantly increased with exposure to HK PeWT at 120 hours (p<0.01), with no detectable effect on flies fed with HK PeGacA-supplemented food (p>0.05). Differences between groups were estimated by post hoc comparisons (Tukey’s honest significant differences) and are indicated by different letters in each plot (p<0.05).

We dissected and stained the guts of three to five-day-old males at three timepoints: before exposure (Untreated), at the end of exposure (72 hours), and 48 hours after transfer to normal food (120 hours) (Fig. 4B).

Quantification of number of positive cells for apoptotic marker Dcp-1 revealed that exposure to HK PeGacA is sufficient to induce an increase in apoptosis at 72 hours (EMMs(zinf): est= -3.9 (SE= 1.0), p= 0.003), which resolves after removal from the treatment food and into clean food 48 hours later (EMMs(zinf): est= -0.6 (SE= 0.7), p= 1) (Fig. 4C, Fig. S4, Tables S1, S7). Additionally, quantification of number of positive cells for mitotic marker PH3 detected no significant differences at either timepoint for this treatment (Fig. 4C, Fig. S4, Table S7). However, exposure to HK PeWT led to a major increase in the number of apoptotic cells at 72 hours of feeding (EMMs(zinf): est= -36.0 (SE= 4.2), p= 6.5E-14). This number remained elevated 48 hours after treatment removal and exposure to clean food, although the effect was not statistically significant (EMMs(zinf): est= -7.2 (SE= 2.5), p= 0.06). This trend supports the deleterious effect of HK PeWT on gut-cell integrity. In line with this, we observed a significant increase in the number of dividing cells (EMMs(zinf): est= - 10.9 (SE= 3.1), p= 0.01) for HK PeWT guts at 120 hours, suggesting a process of gut health restoration and cell renewal in flies recovering from damage (Fig. 4C, Fig. S4, Table S7).

Together, these results show that exposure to heat-killed *P. entomophila* causes tissue damage and underscores the role of tissue repair response against the action of bacterial virulence factors.

### 5. The effects of exposure to heat-killed bacteria are influenced by the microbiota

Further, we tested the role of the gut microbiota in the mortality phenotype observed upon exposure to heat-killed *P. entomophila.* For that, we exposed germ-free and non-germ-free flies to HK PeWT or HK PeGacA and subsequently maintained them on either germ-free or regular food (Fig. 5). With this design we were able to separately assess the role of the microbiota prior to and after exposure to heat-killed bacteria.

**Figure 5.**
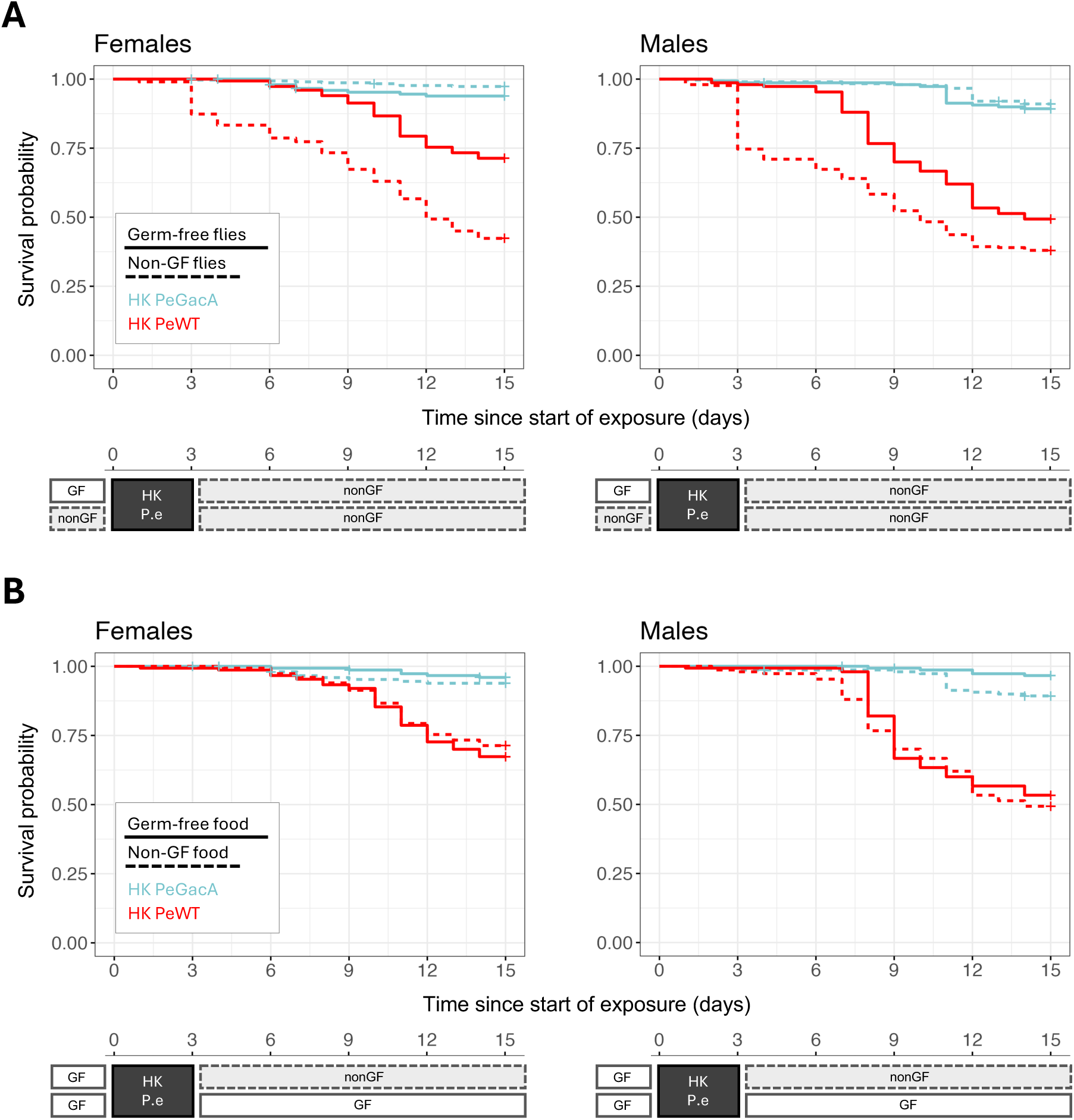
Effects of microbiota on the response to HK *P. entomophila* exposure. Survival curves followed for 14 days for females (left panels) and males (right panels) exposed to heat-killed (HK) *P. entomophila* wildtype (HK PeWT in red) or ΔGacA mutant (HK PeGacA in light blue)**. A)** Survival curves of germ free (solid lines) and non-germ free (dashed lines) flies. Survival differed between germ free and non-free flies when exposed to HK PeWT in both sexes (p<0.001) but not when exposed to HK PeGacA (p>0.05). Flies exposed to HK PeGacA show reduced mortality compared to HK PeWT flies, independently of the germ-free treatment (p< 0.05). **B)** Survival curves of flies transferred to germ free food (solid lines) or normal food (dashed lines). Once again, flies exposed to HK PeWT show higher mortality than those exposed to HK PeGacA (p<0.001), but no differences were found in flies maintained in germ-free *vs*. normal food (p> 0.05 in both sexes) (Tables S1, S8).

As before, we found differences in survival between flies exposed to HK PeWT or HK PeGacA (Anova(Cox): χ²(1)= 18.8, p= 1.45e-05) and between sexes (Anova(Cox): χ²(1)= 8.15, p= 0.004) (Fig. 5, Table S1). We also found that the response to HK WT, but not GacA *P. entomophila*, differed between germ free and non-germ free animals (Anova(Cox): χ²(1)= 12.05, p= 5e-04) (Fig. 5A). Mortality was significantly lower in germ-free than in non-germ-free flies exposed to HK PeWT (EMMs(Cox): p<0.001 for both sexes). This reduction in survival in the presence of microbiota was specific to HK PeWT and not seen with HK PeGacA (Tables S1, S8). Finally, we found no significant effect of the type of food flies were reared on after exposure to HK bacteria (Anova(Cox): χ²(1)= 0.32, p= 0.57) (Fig. 5B, Tables S1, S8).

### 6. Exposure to heat-killed *P. entomophila* triggers an immune response in *D. melanogaster*

We next sought to test how the host immune response reacted to exposure to heat-killed *P. entomophila* (Liehl et al., 2006; Vodovar et al., 2005) and measured expression levels of canonical immunity and stress pathway genes. For that, we exposed flies to HK PeWT or HK PeGacA and extracted RNA from adult males, either untreated or at three different time points post-exposure, 8h, 72h, or 96h.

We measured the transcriptional profile of canonical antimicrobial peptides (AMPs), reactive oxygen species (ROS), and stress response pathway genes (De Gregorio et al., 2002; Hoffmann, 2003; Lemaitre & Hoffmann, 2007). In particular, we selected genes from the JAK-STAT stress response pathway (i.e. transcription factor *STAT-92E*, its cytokine Unpaired-3 (*Upd3*)) and two of its gut effectors, Drosomycin-like 2 (*Drsl-2*) and Drosomycin-like 5 (*Drsl-5*)). We included genes from both Toll and the IMD pathways (i.e. Diptericin (*Dpt*), Attacin-A (*AttA*) for IMD, Drosomycin (*Drs*) for Toll, and Cecropin (*Cec A*) for both), as well as from the ROS pathway (i.e. Dual-oxidase (*Duox*)) (Buchon, Broderick, Chakrabarti, et al., 2009; De Gregorio et al., 2002; Ferrandon et al., 1998; Tzou et al., 2000) (Fig. 6).

**Figure 6.**
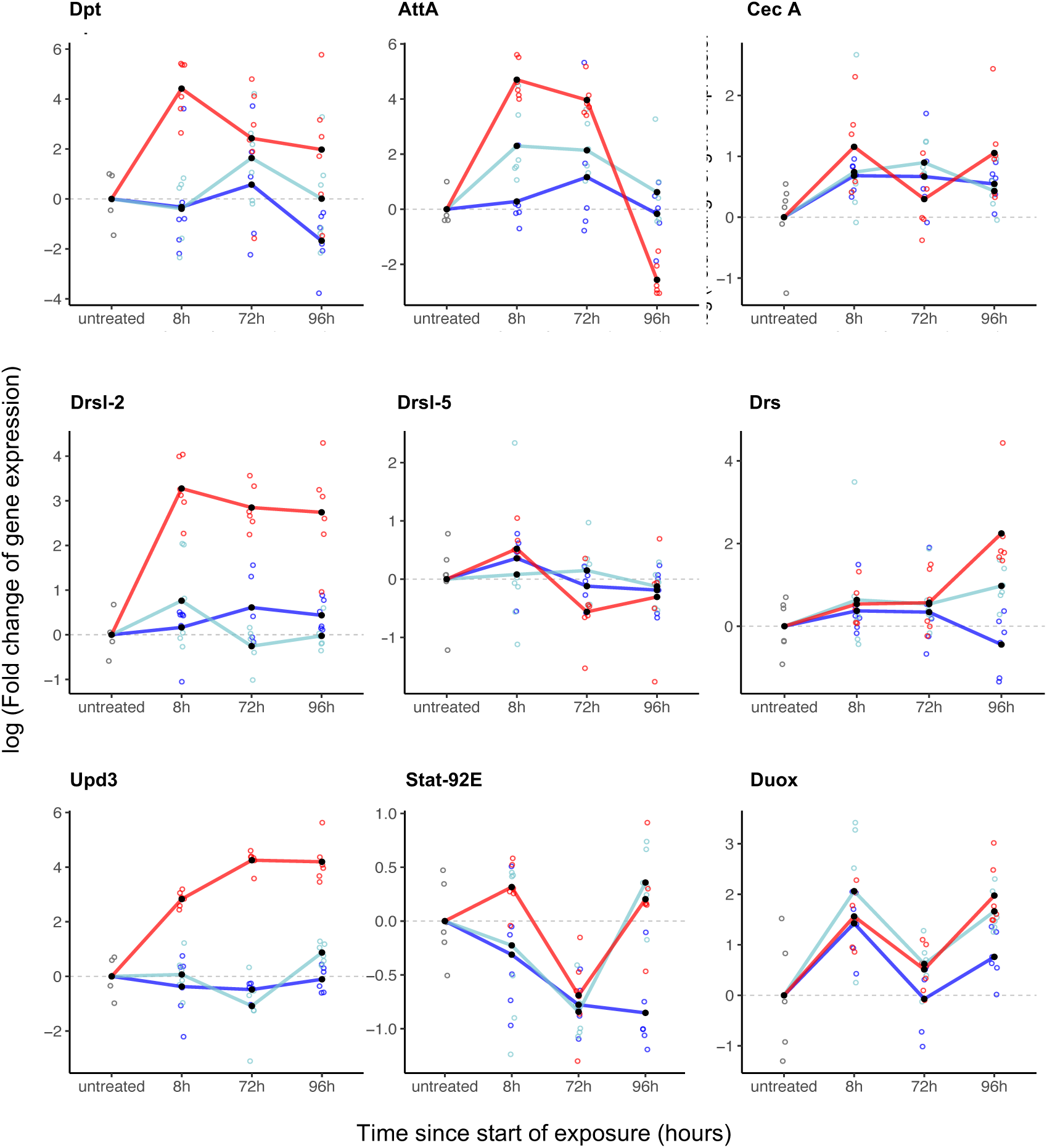
Exposure to HK *P. entomophila* triggers an immune response in *D. melanogaster*. Gene expression (represented as log10 fold change expression normalized to the housekeeping gene Eif-2) in cDNA from head-less males previously exposed to HK PeWT (red), HK PeGacA (light blue), or PBS control (dark blue). Samples were collected at three different timepoints: 8h, 72h, and 96h after exposure, as well as from untreated individuals. qPCR was performed for nine canonical immune genes.

Overall, we observed that several genes were upregulated in flies exposed to HK PeWT as compared to those exposed to either PBS or HK PeGacA (Fig. 6, Tables S1, S9). None of the six AMPs showed noticeable expression differences throughout time under the PBS or HK PeGacA treatments (EMMs(lm) with p>0.05, Table S9). The same unresponsiveness was observed for three AMPs in the HK PeWT treatment, namely *Drsl-5, Cec A*, and *Drs* (EMMs(lm) with p>0.05, Table S9) (Fig. 6). In contrast, three other AMPs, *Drsl-2, Dpt*, and *AttA,* were significantly upregulated within the first eight hours of HK PeWT exposure treatments (EMMs(lm) with p<0.05, Table S9). Interestingly, *Upd3* showed a similar expression pattern to *Drsl2*, with a consistent up-regulation across all timepoints, exclusively under HK PeWT (EMMs(lm) with p<0.05 for all timepoints, Table S9). *STAT-92E*, on the other hand, showed a specific delayed upregulation pattern, detectable at 96h under both HK PeWT (EMMs(lm): est= -1.1 (SE= 0.2), p= 0.03) and HK PeGacA (EMMs(lm): est= -1.2 (SE= 0.2), p= 0.0003) treatments. Finally, *Duox* was constant and unchanged throughout time points and across treatments (Fig. 6, Table S9).

With these results, we conclude that exposure to HK *P. entomophila* triggers stress and immune response pathways in *D. melanogaster*, in a time- and virulence factor-dependent manner

## Discussion

In this study, we sought to measure disease tolerance in *Drosophila melanogaster* upon oral exposure to *Pseudomonas entomophila* by using an inactivated form of the bacterium, thus fixing host resistance mechanisms. We reasoned that orally exposing *Drosophila* to heat-killed bacteria would trigger its immune response pathways through the detection of antigens that withstood the inactivation treatment. These include cell wall components, such as DAP-type peptidoglycans found in gram-negative bacteria, virulence factors (and toxins in the case of *P. entomophila*), and other pathogen-associated molecular patterns (PAMPS) (Buchon et al., 2014; Hoffmann, 2003; Troha et al., 2018; Vincent & Dionne, 2021; Wen et al., 2019). Bacterial heat inactivation has been successfully used for decades, particularly in the mouse model, to trigger an inflammatory response (Jung et al., 2017; Kawase et al., 2012; Minute et al., 2024; Whitlock et al., 2008), but also in insect species such as in the silkworm *Bombyx mori* (Miyashita et al., 2014, 2015). *Pseudomonas entomophila* is particularly suitable for this procedure because it is a natural pathogen of *Drosophila* that secretes toxins, inducing strong host responses (Acken & Li, 2023; Dieppois et al., 2015; Nonaka et al., 2020; Vallet-Gely et al., 2010; Vodovar et al., 2005). Additionally, as a natural pathogen of *D. melanogaster*, it is likely to more faithfully replicate in the laboratory, ecologically relevant host defense processes.

Feeding flies for 24 hours with live *P. entomophila* negatively affects survival, with observable effects after six hours of continuous feeding (Dieppois et al., 2015; Neyen et al., 2014; Vallet-Gely et al., 2010). In contrast, our observations (unpublished data) and other reported studies (Vincent & Dionne, 2021), show that 24 hours of feeding on HK bacteria does not increase mortality. This informed the development of our feeding protocol by exposing flies to HK bacteria for longer periods, revealing a sustained and repeatable effect on host survival and reproduction (Fig. 1). In this context, due to the different duration of exposures, directly comparing mortality caused by live infection against HK bacteria treatment is not strictly appropriate. However, analyzing these differences remains useful and informative for identifying the recovery mechanisms underlying disease tolerance. By contrasting two *P. entomophila* strains - the wild-type and the non-virulent ΔGacA mutant (Fig. 3) (Vallet-Gely et al., 2010; Vodovar et al., 2005, 2006) - we attribute a causal role to virulence factors in driving reduced survival upon oral exposure to heat-killed *P. entomophila*. Within this experimental framework, we show that the presence and activity of virulence factors are pivotal in eliciting disease tolerance responses, as exposure to heat-killed bacteria mirrors the fitness consequences of a live infection (Figs. 2, 3).

### 1. *P. entomophila* virulence factors activate host immune response

Activation of the *Drosophila* immune response has been associated to shortened life span (Fabian et al., 2018; Garschall & Flatt, 2018; S. O. Zhou et al., 2024), changes in metabolism (most notably in lipid, carbohydrate, and protein metabolism) (Bland, 2023; Cumnock et al., 2018; DiAngelo et al., 2009; Dionne, 2014; Loudhaief et al., 2023; Socha et al., 2023) and neurodegeneration (Nayak & Mishra, 2022). We show in this study that the presence of virulence factors correlates with immune activation, quantified by the expression of effector genes including anti-microbial peptides (AMPs), oxidative stress, and inflammation mediators (Fig. 6).

We characterized and compared the expression levels of nine genes at four specific time points: 0h (untreated flies), 8h upon exposure, 72h (end of exposure) and 96h (24h post-exposure), which we anticipated would provide insights onto the recovery phase following exposure. Gene expression level dynamics were, in most cases, different between the wild-type strain, and the avirulent ΔGacA mutant or PBS controls, underscoring the link between bacterial virulence and the activation of immune responses (Diard & Hardt, 2017; Gates et al., 2021; Horrocks et al., 2011).

Albeit at different stages or routes of infection, previous reports have shown that live *P. entomophila* triggers strong changes in the expression profiles of more than 200 genes including AMPs such as Diptericin, Drosomycin, Cecropins, and Attacins (Buchon, Broderick, Poidevin, et al., 2009; Troha et al., 2018; Vodovar et al., 2005). Consistently with such previous observations, we found that feeding on HK PeWT bacteria promotes the rapid (within 8 hours) expression of three out of six AMPs (*AttA, Dpt,* and *Drsl-2*), while *Drs* showed an upregulation only in the recovery period, i.e. 24 hours after the end of exposure (96h timepoint). This unresponsiveness of *Drs* has been observed under oral infection in larvae (Vodovar et al., 2005), but not in other contexts namely in adult systemic infections (Troha et al., 2018) or in gut microbiota control during metamorphosis (Nunes et al., 2021), where it is upregulated. Together, these data reinforce the context-dependent role of AMPs and their intricate upstream regulation (Hanson et al., 2019; Hanson & Lemaitre, 2020; Vincent et al., 2022), underscoring the need for a systematic analysis of the underlying regulatory processes that incorporates diverse life-history traits and their ecological relevance.

In this context, as with other epithelial tissues, these three AMPs are effectors of the IMD and/or JAK/STAT pathways (Buchon et al., 2014; Hoffmann, 2003; Hultmark, 2003; Myllymäki & Rämet, 2014). Accordingly, *Upd3*, another JAK/STAT pathway member, shows sustained upregulation upon HK bacteria exposure. Damage to the gut epithelium has been well documented upon oral live *P. entomophila* infection, with mortality being primarily attributed to the pathogen and the action of its virulence factors, which inhibit host translation and impair cell repair processes. (Chakrabarti et al., 2012; Jiang et al., 2009; Vodovar et al., 2005). Such damage is known to activate the JNK and Hippo pathways, which in turn leads to the activation of the JAK-STAT pathway via the cytokine *Upd3* to promote the proliferation of intestinal stem cells, necessary for tissue repair (Buchon, Broderick, Poidevin, et al., 2009; Buchon et al., 2010; Myllymäki & Rämet, 2014; F. Zhou et al., 2013). Given the above, it is reasonable to speculate that the presence of HK *P. entomophila* and its virulence factors, which cause epithelial damage (Fig. 4), closely mimic the effect of live infections, thereby triggering the JAK/STAT-mediated repair process.

The production of ROS is a hallmark of live pathogenic infections, notably in *P. entomophila* infections (Bae et al., 2010; Buchon, Broderick, Poidevin, et al., 2009; Chakrabarti et al., 2012; Ha et al., 2005; Zeng et al., 2022). However, our results fail to show evidence of ROS activity, as there was no upregulation of *Duox* at any of the time points tested. Nonetheless, it is difficult to completely rule out the involvement of oxidative stress in the reported phenotypic effects. Several possible, but not mutually exclusive, explanations could account for the lack of *Duox* transcriptional response. Firstly, the transcriptional unresponsiveness of *Duox* may be attributed to our experimental design, which may not have captured the optimal time window for detecting significant changes in expression. While *Duox* is crucial for the pathogenicity of live *P. entomophila* oral infection, no significant transcriptional differences are observed in fly guts as early as 4h post-infection (Buchon, Broderick, Poidevin, et al., 2009; Chakrabarti et al., 2012). Secondly, a more direct measurement of the volatile and transient oxidative-stress effector molecules, such as assessing cytochrome c reduction levels, may be necessary to accurately evaluate the role of ROS in the fitness costs induced by ingesting HK bacteria (Murphy et al 2022). Thirdly, the unchanged transcriptional profile of *Duox* could be explained by the lack of Uracil production in our heat-killed bacterial context. Uracil has been shown to play a key role in helping the host distinguish between commensals and pathogens, triggering various elimination mechanisms (Kuraishi et al., 2013; Lee et al., 2013). In our setup, heat-killed bacteria likely provide a level of Uracil that is, at best constant, and probably decreasing over time, which may explain the negative result.

Future experiments should consider these factors into account to more directly investigate the role of oxidative stress. This will be a crucial step in further elucidating the mechanisms that cause damage upon exposure to HK bacteria.

### 2. Heat-killed bacteria induces gut damage and repair in a virulence factor-dependent way

We demonstrated that exposure to a virulent strain of *P. entomophila* induces significantly higher levels of cell death in the gut epithelium than its non-virulent counterpart, the ΔGacA mutant (Fig. 4, Fig. S3). This confirmed our hypothesis that oral exposure to heat-killed virulent bacteria induces damage, as previously reported for live infections (Buchon, Broderick, Chakrabarti, et al., 2009; Buchon et al., 2013).

One important distinction, however, may lie in the duration of the challenge and the timing of damage and repair. In our experimental framework, exposure to HK PeWT leads to a significant increase in cell death, peaking at the end of the exposure 8- to 10-fold higher than PBS and HK PeGacA treatments (Fig. 4C, left panel). This difference diminishes within 48 hours (at the 120h timepoint), suggesting a temporal limit on the deleterious effects of virulence factor exposure to gut integrity.

This observation aligns with the upregulation of the previously described responses involving JAK/STAT activation and concomitant *Upd3*-mediated increase in expression (Fig. 6), shown to promote enterocyte renewal and restoration of epithelium homeostasis (Kuraishi et al., 2011; F. Zhou et al., 2013). This upregulation is initiated within the first hours of exposure, persists throughout the exposure period, and continues for at least 24 hours after the challenge has ended.

In contrast, the expression profile of the JAK/STAT pathway transcription factor *Stat-92E* exhibits more complex dynamics. It is initially upregulated within 8 hours of exposure, then downregulated to homeostatic levels (as defined by PBS treatment) by 72h, the peak of exposure time. This is followed by a significant increase 24 hours later, upon transfer to clean food (Fig. 6). Evidence suggests that although there is upregulation of damage cytokines (such as *Upd3*) that trigger downstream pathways leading to epithelial repair, this response is blocked due to a translational arrest caused in part by the bacterial toxin Monalysin (Chakrabarti et al., 2012; Nonaka et al., 2020; Opota et al., 2011). This effect could explain why at 72h there is no noticeable difference in cell proliferation between treatments (Fig. 4C, right panel). However, the number of proliferating cells at 120h, i.e. 48 hours upon release from exposure and presumably the clearance of Monalysin, increases significantly in syntony with the second wave of *Stat-92E* upregulation (Fig. 6).

Altogether, these observations support previous evidence of a pathogen-dependent impairment in epithelial repair, which likely plays a role in the death of the fly (Buchon, Broderick, Chakrabarti, et al., 2009).

### 3. Microbiota presence lowers the survival of HK-exposed flies

Microbiota profoundly influences host health and physiology, including digestive and immune functions (Bäckhed & Crawford, 2010; Broderick & Lemaitre, 2012; Round & Mazmanian, 2009). In *Drosophila*, reports show that certain aspects of the microbiota can improve survival and longevity. However, the relationship is complex and context-dependent (Hong et al., 2022; Tafesh-Edwards & Eleftherianos, 2023). In our experimental framework, mortality following exposure to HK PeWT is exacerbated in the presence of the microbiota, while exposure to HK PeGacA does not show this effect (Fig. 5), reiterating the pivotal role of virulence factors in influencing the observed outcomes.

Interestingly, this deleterious effect of the microbiota in survival to HK PeWT feeding depends on when it is present relative to the exposure itself. Flies exposed to HK PeWT with microbiota present throughout the experiment (dashed red lines in Fig. 5A), show higher mortality compared to those that were germ-free until being transferred to regular food, post-exposure (solid red lines in Fig. 5A). In essence, this difference is driven by whether the microbiota was present or not before and during exposure to HK PeWT. This interpretation is supported by the conditions tested in Fig. 5B, where the survival difference disappears when flies are germ-free prior to exposure, regardless of their subsequent microbiota status.

We can propose two hypotheses to accommodate these observations. Possibly, exposure to the wild-type HK bacteria causes dysbiosis, leading to a compound effect of tissue damage and systemic inflammation, which increases host mortality, as described in previous studies (Jones et al., 2013; Ryu and Kim et al., 2008; Sartor, 2008). Alternatively or concomitantly, increased mortality in flies with microbiota may result from a shift from localized to systemic response. This would be facilitated by the epithelial damage caused by virulent wild-type bacteria and, possibly, peritrophic matrix deterioration (Kuraishi et al., 2011) or from signaling to the fat body upon bacterial peptidoglycan detection (Neyen et al., 2012).

Further work is required to test these hypotheses, particularly to understand if and how localized gut damage from ingesting virulent bacteria translates into a systemic response or how the presence of gut microbiota impacts the expression of important immunity effector genes, in the context of the HK PeWT exposure.

### 4. Pervasive sexual dimorphism in survival upon exposure to HK *P. entomophila*

In *Drosophila* studies, sexual dimorphism has been reported in several aspects of fly physiology including in their immune response (Belmonte et al., 2020; Shahrestani et al., 2018). Studies with *Drosophila* have shown that the sexes exhibit profound differences in survival and pathology in response to infections with different kinds of pathogens, including bacteria and viruses. Males are known to be more susceptible than females to some acute viral infections (Gupta, Stewart, et al., 2017; Palmer et al., 2018), while the opposite is true in bacterial and fungal infections (Belmonte et al., 2020). It is important to note, however, that very few studies utilize both sexes for infection-based studies in *Drosophila*. In addition to this, survival outcomes after infection are heavily dependent on the nature of pathogen and on other external factors like host diet and age (Bashir-Tanoli & Tinsley, 2014; Le Bourg, 2011; Parker et al., 2020). Interestingly, male flies consistently show higher mortality in response to HK *P. entomophila* exposure across all our experiments (Figs. 1, 2, 3 and 5), the opposite to what is typically observed upon oral bacterial infection (Belmonte et al., 2020; Duneau et al., 2017).

The higher mortality observed in males after exposure to HK *P. entomophila* could be explained by different energy allocation to immunity between sexes, as shown by classical examples of immunity/reproduction trade-offs (Chang van Oordt et al., 2022; French et al., 2007; Luckinbill et al., 1984; Schwenke et al., 2016; Syed et al., 2020; Villafranca et al., 2023) and suggested by a faster activation and more prolonged immune response reported in *D. melanogaster* males (Vincent & Dionne, 2021).

Another possibility explaining the sexual dimorphism we observe, relies on described sex differences in intestinal physiology (Hudry et al., 2016; Regan et al., 2016; Reiff et al., 2015), notably in the proliferation of intestinal stem cells (ISC) (Lemaitre & Miguel-Aliaga, 2013; Miguel-Aliaga et al., 2018). Indeed, females repair gut epithelial damage at much faster rates compared to males upon infection or detergent-induced gut damage (Regan et al., 2016, 2022). Recent evidence also shows that epithelial growth factor receptor (EGFR) mutant males display a severe loss of disease tolerance as compared to mutant females (Prakash et al., 2022). Alternative hypotheses for the observed dimorphic survival outcomes—seemingly more prosaic—might include differences in body size, which can influence metabolic rates, immune function, and overall susceptibility to infection (Teder & Tammaru, 2005). Larger body size in females, for instance, could confer some level of advantage in resource allocation for immune responses, making them more resilient to the same pathogen. Investigating both genetic and ecological factors that contribute to these sex-specific outcomes will be important for a deeper understanding of disease tolerance and survival strategies across sexes.

### 6. Exposure to heat-killed *P. entomophila* entails fecundity costs

Alongside the extensive characterization discussed above on the effects of exposure to heat-killed *P. entomophila* on survival, we have explored the impact on the other core fitness component, reproduction. Infection with HK bacteria in *Drosophila* is known to result in decreased fecundity (Bashir-Tanoli & Tinsley, 2014). This reduction in fecundity, comparable to that caused by live bacteria, had also been reported and attributed to the cost of activating an immune response (Zerofsky et al., 2005). We have replicated these observations with both heat-killed and live bacteria (Fig. 3B, Fig. S3B). Given that exposure to HK bacteria triggers an immune response (Troha et al., 2018; Wen et al., 2019), the reduction in fecundity could be explained in part by the immunity-reproduction trade-off previously reported in *Drosophila* and other insect models, as mentioned above (Chang van Oordt et al., 2022; French et al., 2007; Luckinbill et al., 1984; Schwenke et al., 2016; Syed et al., 2020; Villafranca et al., 2023). Immune response and reproduction are both physiologically and energetically demanding processes, requiring the allocation of substantial resources, either from stored reserves or through reallocation (Bashir-Tanoli & Tinsley, 2014). Ultimately, prolonged immune activation may lead to energy depletion, which could be reflected in other fitness traits, such as fertility and fecundity (Meiselman et al., 2018).

Additionally, we also observe that the fecundity reduction correlates strongly with bacterial virulence and not with pathogen species *per se*, as our experiments with other non- or mildly-virulent gram-negative bacteria species revealed a higher reproductive output as compared to the HK PeWT-exposed group (Fig. 2). This was confirmed after observing increased fecundity in flies exposed to the avirulent form of the *P. entomophila* under both live and HK conditions (Fig. 3B, Fig. S3B). This effect is supported by the consistent increase in fecundity observed in females fed any heat-killed bacteria diet. This difference can be attributed to the nutritional quality of the diet whereby the consumption of non-virulent HK bacteria has a greater nutritional value, positively impacting reproductive investment (Keebaugh et al., 2018). In contrast, consuming heat-killed bacteria containing virulence factors would impose significant costs on the host, through tissue damage, that the nutritional benefit of the food cannot compensate.

Our results reveal variation in fecundity among flies surviving heat-killed bacterial exposure, suggesting an evolutionary trade-off hinging on disease tolerance. It has been proposed that variation in disease tolerance between individuals (genotypes) may result from a distinct balance between tissue repair and reproduction (Kutzer et al., 2016, 2018; Syed et al. 2020). This finding highlights the need for further research exploring the genetic and physiological mechanisms driving disease tolerance for which our experimental framework may contribute.

This experimental framework has enabled us to integrate fitness traits (i.e., survival and reproduction) to clarify the relationship between host immune response activation and its potential fitness cost. This approach can not only enhance our understanding of how hosts reduce fitness costs through disease tolerance mechanisms but also provide insights into how evolution may shape immune strategies against pathogens. While our findings are specific to *P. entomophila* and oral infection, future studies will examine how disease tolerance varies across different pathogens and infection routes, offering deeper insights into host-pathogen interactions and the interplay between disease tolerance and resistance mechanisms.

## Materials and Methods

### Fly stocks and rearing conditions

An outbred *Drosophila melanogaster* population was established and maintained in the lab since 2007 (Faria et al., 2015; Martins et al., 2013, 2014; Paulo et al., 2023). This population henceforth referred to as ‘Mel2’ was used for all experiments unless otherwise stated. The population is maintained in laboratory cages at a census of 1500-2000 individuals, under constant temperature (25°C) and humidity (60-70%) with a 12:12 light-darkness cycle and fed with cornmeal-agar medium, consisting of 4.5% molasses, 7.5% sugar, 7% corn-flower, 2% granulated yeast extract, 1% agar, and 0.25% nipagin, mixed in distilled water. All experimental flies were generated from egg lays with controlled density. Three- to five-day-old, mated flies were used for all experiments.

### Bacterial strains and culture

All bacterial strains were cultured in an LB medium with the appropriate antibiotics, when applicable. *Escherichia coli* K-12 (*E. coli*-K12) was grown at 37°C according to standard protocols. *Erwinia carotovora carotovora 15* (Ecc-15), *Pseudomonas entomophila,* and its mutant strain ΔGacA (Opota et al., 2011; Vallet-Gely et al., 2010; Vodovar et al., 2006), were kindly shared by the Lemaitre Lab at the École Polytechnique Fédérale de Lausanne, as well as *P. putida* from lab generated stocks were grown at 29°C according to established protocols (Faria et al., 2015; Martins et al., 2013). Briefly, frozen stocks of bacteria were plated to develop colonies overnight. A single colony was then streaked to make a 5ml starter culture under aseptic conditions and grown in an orbital shaker at 180 rpm for 6 hours. Overnight cultures were then made with a larger volume of Luria Bertani (LB) media using the confluent starter cultures. For live infections, bacteria pellets were collected after centrifugation at 4600rpm for 15 minutes at 4°C and resuspended in sterile LB. For treatment under heat-killed conditions, bacterial pellets were resuspended in sterile PBS and adjusted to the desired concentration (OD600= 100 for *P. putida*, *Ecc-15,* as well as for all strains of *P. entomophila*, and OD600= 200 for *E. coli*).

For the bacteria inactivation with heat, resuspended bacteria with the desired OD were heat-killed for one hour in a water bath at 55°C, for *P. entomophila* and its mutant strains as well as for the two *Ecc* strains, or at 70°C for *E. coli*. These were the minimal temperatures that did not exhibit any growth (CFUs) upon subsequent plating and overnight incubation at 29°C. Bacteria inactivation using the fixative paraformaldehyde (PFA) was confirmed in the same manner and prepared by incubating resuspended bacteria pellets in 1% PFA for 30 min. The inactivated bacteria were subsequently washed three times with PBS by centrifugation at 4600 rpm for 10 min at each washing step.

### Fly food preparation and exposure to inactivated bacteria

Liquid fly food was prepared using standard laboratory conditions (as described previously) and mixed with the inactivated bacteria in a ratio of 1:1. To account for the texture and viscosity of the food, extra agar powder was added to the mixture (to give an overall 100% agar content after dilution of bacteria) before the addition of the required volume of inactive bacteria solution. In the end, food mixed with agar was dispensed into fly bottles and left to cool down and solidify for at least two hours before use. As a control for the treatment food with bacteria, all the steps were repeated but using sterile PBS instead of inactive bacteria.

### Germ-free *Drosophila* experiment

We set egg lays for 1-hour period in agar plates. Collected eggs were washed with distilled water, transferred to 2% bleach for 10-15min, washed twice in distilled water, transferred to 70% ETOH for 5min, washed twice in distilled water. We transferred 30-50 eggs to vials with sterile food (autoclaved) and left in sterile tray until eclosion. Under sterile conditions we generated five replicate vials each containing 10 males and 10 females in either sterile food or regular food in which laboratory flies had been kept on for 24 hours (hence, inoculating the medium with their microbiota). Mortality was followed daily for 15 days with regular flipping to new vials under the same conditions.

### Infection with live bacteria

Oral infection of *D. melanogaster* with *P. entomophila* was performed using the previously established protocols (Martins et al., 2013; Paulo et al., 2023). Briefly, starter and overnight cultures were grown using sterile LB media at 29°C as previously described. After adjusting the concentration of the pelleted cells to the desired concentration (OD600= 100) they were diluted 1:1 with 5% sucrose solution, prepared under sterile conditions just before the infection. The resulting mixture was pipetted into filter papers and placed inside food vials to which we transferred groups of 10 single-sex flies to feed for 24h at 25°. Later, they were moved to clean vials with normal food where survival was measured for 6 days. As a control to the infection experiment, flies were fed with the 5% sucrose solution without the bacteria.

### Phenotyping survival and fecundity

Flies were mated and aged between three to five days for all experiments. Bacteria exposure treatment was done with 15 females and 15 males per vial except for experiments with bacterial mutants, where we used 10 males and 10 females per vial. Survival measurements began at the start of the treatment and lasted 21 days or 14 days, depending on the experiment (see Results). Flies that escaped or sustained accidental injury during transfer were censored from the analysis, as well as flies that died within the first hour of the experiment.

Fecundity was measured by transferring adults daily to new vials and counting the number of pupae produced each day (Fig. S1). We have tested pupal to adult viability in our experimental conditions, to find that it is close to 100%, validating pupae count as a strong proxy for fecundity (data not shown). Melanized Pupae or third-instar larvae were excluded from the count. Fecundity was calculated by normalizing the number of pupae to the daily count of live females, providing an estimate of daily fecundity per female. Lifetime fecundity per female was determined by summing these daily values over the specified period.

### Gut sample staining and quantification

Intestines of three-to-five-day-old males exposed to the different HK bacteria treatments at specific time points were dissected in cold 1xPBS and immediately fixed in 4% formaldehyde for 30 mins. Fixation followed three 15-minute washes with PBS-T (0.1% Triton X-100/PBS) and blocking for 1 hour at room temperature with 2% BSA (2% BSA/0.1% Triton X-100/PBS). Guts were then incubated overnight at 4° C with primary antibodies diluted in blocking solution (1:200 rabbit anti-Dcp1, Cell Signaling #9578; 1:500 mouse anti-PH3, Cell Signaling #9706) followed by three washes in PBS-T and a 2-hour room temperature incubation with secondary antibodies (anti-rabbit Alexa Fluor 568; anti-mouse Alex Fluor 488) and (1:1000) DAPI. After three washes, samples were mounted in 90% glycerol. Images were acquired on a Leica TCS SP5 confocal microscope, using a 20x/0.7 objective and 405, 488, and 532 laser lines to excite DAPI, Alexa488, and Alexa568, respectively. Images of ∼6 male midguts were used to manually quantify fluorescent cells for Dcp-1 (a marker for apoptosis; REF) and PH3 (a marker for proliferation; REF) over an area normalized by the number of total cells (DAPI).

### RNA extractions and RT-qPCR

For all RNA extractions, flies were exposed to the protocol described above and, at different time points (0h or untreated, 8h, 72h, and 96h post-exposure) three males per treatment per time point were pooled with their heads removed, and subsequently manually homogenized using a sterile pestle in 500 µL of Trizol. RNA extractions were performed using a phenol-chloroform protocol from which 1 µg of RNA were used per pooled sample replicate for cDNA synthesis. After precipitation, a DNase I (RQ1 RNASE-FREE DNASE 1* from Promega®) treatment was done to all samples, followed by reverse transcription using the Thermo Scientific® RevertAid H Minus cDNA kit. cDNA was finally diluted 1:5 for the qPCR. For quantification of gene expression, qPCRs were performed using SYBR^TM^ Green Master Mix (Thermo Scientific®) and reactions ran on 384-well plates (Applied Biosystems®). The PCR conditions used in all experiments were: initial denaturation/ enzyme activation, 95°C for 10’; followed by 45 cycles of denaturation, 95°C for 10”; annealing, 60°C for 10”; extension, 72°C for 30”. Sequences of primers used for qPCRs are shown in Supp. Table. 27 with *EIF2* as a reference housekeeping gene. Gene expression analysis was performed using relative quantification (ΔΔCt), consisting of taking the average of the technical replicates for each candidate gene and subtracting the average of the Ct values of the respective sample’s housekeeping gene (*EIF2*) (ΔCt) and normalizing this value to the ΔCt of the respective reference condition.

### Statistical analysis

All statistical analyses were conducted in R v4.2.1 (R Core Team, 2021) using the following R packages: *tidyverse* (Wickham et al., 2019) for data manipulation, *ggplot2* (Wickham, 2016) for visualization, *lme4* (Bates et al., 2015) and *lmerTest* (Kuznetsova et al., 2017) for linear mixed-effects models, *glmmTMB* (Brooks et al., 2017) and *pscl* (Zeileis et al., 2008) for zero-inflated models, and *emmeans* (Lenth, 2017) for post-hoc multiple comparisons.

Survival data were analyzed using Cox proportional hazards models, followed by Type III ANOVA to assess the effects of fixed factors. Fecundity data (measured as the sum of daily fecundity) and data on the number of Dcp-1 or PH3 positive cells were analyzed using zero-inflated negative binomial models. Exceptionally, for the experiment in Fig.1, fecundity data was measured as the number of pupae by the end of the experiment, and this data was analyzed using linear mixed-effects models. For both survival and fecundity data, post-hoc pairwise comparisons among treatment groups were conducted using estimated marginal means, with significance set at p < 0.05. For all experiments, models included relevant fixed factors (e.g., Treatment, Sex, Days of Exposure, Bacterial species, Bacterial strain), while accounting for random effects from bottles or vials where applicable. Full model details and outputs are provided in the supplementary tables.

## Acknowledgements

We are indebted to Caren Norden and her lab for critical logistical support. We thank Bruno Lemaitre, Isabel Gordo and Luis Teixeira for reagents and stocks, Liliana Vieira and Sandra Crisóstomo for technical assistance, André Barros for help with analyses, and Diogo Roque, David Duneau, João Picão-Osório, Patrícia Duarte, Ricardo Vieira, and Tanvi Madaan, for discussions that improved the work and manuscript. All imaging was performed at the Faculty of Sciences of the University of Lisbon’s Microscopy Facility, which is a node of the Portuguese Platform of BioImaging, reference PPBI-POCI-01-0145-FEDER-022122. This work was supported by Instituto Gulbenkian de Ciência/Fundação Calouste Gulbenkian and FCT-Fundação para a Ciência e a Tecnologia (Portugal), through UID/Multi/04555/2013 and UIDB/00329/2020. FCT-Fundação para a Ciência e a Tecnologia (Portugal) funded the PhD fellowships of PAA (BD/06404/2020) and of TFP (BD/128432/2017 and COVID/BD/151645/2021), and the project PTDC/BIA-BIO/4693/2021. Support was also provided by C.S. CONGENTO, project LISBOA-01-0145-FEDER-022170, co-financed by Lisboa Regional Operational Programme (Lisboa 2020), under the Portugal 2020 Partnership Agreement, through the European Regional Development Fund (ERDF), and Foundation for Science and Technology (Portugal).

## Author contributions

Conceptualization: Élio Sucena

Data acquisition: Priscilla A. Akyaw, Elvira Lafuente, Tânia F. Paulo

Data Curation: Priscilla A. Akyaw, Elvira Lafuente, Tânia F. Paulo

Formal Analysis: Priscilla A. Akyaw, Tânia F. Paulo, Elvira Lafuente

Funding Acquisition: Élio Sucena

Investigation: Priscilla A. Akyaw, Tânia F. Paulo, Elvira Lafuente, Élio Sucena

Methodology: Priscilla A. Akyaw, Tânia F. Paulo, Elvira Lafuente, Élio Sucena

Project Administration: Priscilla A. Akyaw, Élio Sucena

Resources: Élio Sucena Supervision: Élio Sucena

Validation: Priscilla A. Akyaw, Tânia F. Paulo, Elvira Lafuente

Visualization: Priscilla A. Akyaw, Tânia F. Paulo, Elvira Lafuente

Writing – Original Draft Preparation: Priscilla A. Akyaw, Elvira Lafuente, Élio Sucena

Writing – Review & Editing Preparation: Priscilla A. Akyaw, Tânia F. Paulo, Elvira Lafuente, Élio Sucena

## References

1. Acken, K. A., & Li, B. (2023). *Pseudomonas* virulence factor controls expression of virulence genes in *Pseudomonas entomophila*. PLOS ONE, 18(5), e0284907. 10.1371/JOURNAL.PONE.0284907

2. Acuña Hidalgo, B., Silva, L. M., Franz, M., Regoes, R. R., & O Armitage, S. A. (2022). Decomposing virulence to understand bacterial clearance in persistent infections. Nat Commun, 13, 5023. 10.1038/s41467-022-32118-1

3. Aggarwal, K., & Silverman, N. (2008). Positive and negative regulation of the *Drosophila* immune response. Journal of Biochemistry and Molecular Biology, 41(4), 267–277. 10.5483/bmbrep.2008.41.4.267

4. Arbuthnott, D. (2018). Female life-history trade-offs and the maintenance of genetic variation in *Drosophila melanogaster*. American Naturalist. 192(4), 448–460 10.1086/698727

5. Ayres, J. S. (2017). Microbes Dress for Success: Tolerance or Resistance? Trends in Microbiology, 25(1), 1–3. 10.1016/j.tim.2016.11.006

6. Ayres, J. S. (2020a). Immunometabolism of infections. Nature Reviews. Immunology, 20(2), 79–80. 10.1038/S41577-019-0266-9

7. Ayres, J. S. (2020b). The Biology of Physiological Health. Cell, 181(2), 250–269. 10.1016/J.CELL.2020.03.036

8. Ayres, J. S., Freitag, N., & Schneider, D. S. (2008). Identification of drosophila mutants altering defense of and endurance to *Listeria monocytogenes* infection. Genetics, 178(3), 1807–1815. 10.1534/genetics.107.083782

9. Ayres, J. S., & Schneider, D. S. (2008). A Signaling Protease Required for Melanization in *Drosophila* Affects Resistance and Tolerance of Infections. PLoS Biology, 6(12), e305. 10.1371/journal.pbio.0060305

10. Ayres, J. S., & Schneider, D. S. (2012). Tolerance of Infections. Annual Review of Immunology, 30(1), 271–294. 10.1146/annurev-immunol-020711-075030

11. Bäckhed, F., & Crawford, P. A. (2010). Coordinated regulation of the metabolome and lipidome at the host-microbial interface. Biochimica Et Biophysica Acta, 1801(3), 240–245. 10.1016/j.bbalip.2009.09.009

12. Bae, Y. S., Choi, M. K., & Lee, W.-J. (2010). Dual oxidase in mucosal immunity and host-microbe homeostasis. Trends in Immunology, 31(7), 278–287. 10.1016/j.it.2010.05.003

13. Bashir-Tanoli, S., & Tinsley, M. C. (2014). Immune response costs are associated with changes in resource acquisition and not resource reallocation. Functional Ecology, 28(4), 1011–1019. 10.1111/1365-2435.12236

14. Basset, A. (2000). The phytopathogenic bacteria *Erwinia carotovora* infects *Drosophila* and activates an immune response. Proceedings of the National Academy of Sciences, 97(7), 3376–3381. 10.1073/pnas.070357597

15. Bates, D., Mächler, M., Bolker, B., & Walker, S. (2015). Fitting Linear Mixed-Effects Models Using lme4. Journal of Statistical Software, 67, 1–48. 10.18637/jss.v067.i01

16. Baucom, R. S., & de Roode, J. C. (2011). Ecological immunology and tolerance in plants and animals. Functional Ecology, 25(1), 18–28. 10.1111/j.1365-2435.2010.01742.x

17. Belmonte, R. L., Corbally, M. K., Duneau, D. F., & Regan, J. C. (2020). Sexual Dimorphisms in Innate Immunity and Responses to Infection in *Drosophila melanogaster*. Frontiers in Immunology, 10, 3075. 10.3389/fimmu.2019.03075

18. Best, A., White, A., & Boots, M. (2008). Maintenance of host variation in tolerance to pathogens and parasites. Proc Natl Acad Sci U S A. 105(52):20786–91. www.pnas.org/cgi/content/full/

19. Best, A., White, A., & Boots, M. (2014). The coevolutionary implications of host tolerance. Evolution, 68(5), 1426–1435. 10.1111/evo.12368

20. Bland, M. L. (2023). Regulating metabolism to shape immune function: Lessons from *Drosophila*. Seminars in Cell & Developmental Biology, 138, 128–141. 10.1016/j.semcdb.2022.04.002

21. Bonneaud, C., Tardy, L., Giraudeau, M., Hill, G. E., McGraw, K. J., & Wilson, A. J. (2019). Evolution of both host resistance and tolerance to an emerging bacterial pathogen. Evolution Letters, 3(5), 544–554. 10.1002/evl3.133

22. Brandt, S. M., Dionne, M. S., Khush, R. S., Pham, L. N., Vigdal, T. J., & Schneider, D. S. (2004). Secreted Bacterial Effectors and Host-Produced Eiger/TNF Drive Death in a Salmonella-Infected Fruit Fly. PLoS Biol. 2(12):e418. 10.1371/journal.pbio.0020418

23. Broderick, N. A., & Lemaitre, B. (2012). Gut-associated microbes of *Drosophila melanogaster*. Gut Microbes, 3(4). 10.4161/gmic.19896

24. Brooks, M. E., Kristensen, K., Benthem, K. J. van, Magnusson, A., Berg, C. W., Nielsen, A., Skaug, H. J., Mächler, M., & Bolker, B. M. (2017). glmmTMB Balances Speed and Flexibility Among Packages for Zero-inflated Generalized Linear Mixed Modeling. The R Journal, 9(2), 378–400.

25. Buchon, N., Broderick, N. A., Chakrabarti, S., & Lemaitre, B. (2009). Invasive and indigenous microbiota impact intestinal stem cell activity through multiple pathways in *Drosophila*. Genes & Development, 23(19), 2333–2344. 10.1101/gad.1827009

26. Buchon, N., Broderick, N. A., Kuraishi, T., & Lemaitre, B. (2010). Drosophila EGFR pathway coordinates stem cell proliferation and gut remodeling following infection. BMC Biology, 8(1), 152. 10.1186/1741-7007-8-152

27. Buchon, N., Broderick, N. A., & Lemaitre, B. (2013). Gut homeostasis in a microbial world: Insights from *Drosophila melanogaster*. Nature Reviews Microbiology, 11(9), 615–626. 10.1038/nrmicro3074

28. Buchon, N., Broderick, N. A., Poidevin, M., Pradervand, S., & Lemaitre, B. (2009). *Drosophila* Intestinal Response to Bacterial Infection: Activation of Host Defense and Stem Cell Proliferation. Cell Host and Microbe, 5(2), 200–211. 10.1016/j.chom.2009.01.003

29. Buchon, N., Silverman, N., & Cherry, S. (2014). Immunity in *Drosophila melanogaster*—From microbial recognition to whole-organism physiology. Nature Reviews Immunology 2014 14:12, 14(12), 796–810. 10.1038/nri3763

30. Chakrabarti, S., Liehl, P., Buchon, N., & Lemaitre, B. (2012). Infection-induced host translational blockage inhibits immune responses and epithelial renewal in the *Drosophila* gut. Cell Host & Microbe, 12(1), 60–70. 10.1016/j.chom.2012.06.001

31. Chambers, M. C., & Schneider, D. S. (2012). Balancing resistance and infection tolerance through metabolic means. Proceedings of the National Academy of Sciences of the United States of America, 109(35), 13886–13887. 10.1073/pnas.1211724109

32. Chamy, L. E., Leclerc, V., Caldelari, I., & Reichhart, J.-M. (2008). Sensing of “danger signals” and pathogen-associated molecular patterns defines binary signaling pathways “upstream” of Toll. Nature Immunology, 9(10), 1165–1170. 10.1038/ni.1643

33. Chang van Oordt, D. A., Taff, C. C., Ryan, T. A., & Vitousek, M. N. (2022). Timing of Breeding Reveals a Trade-Off between Immune Investment and Life History in Tree Swallows. Integrative and Comparative Biology, 62(6), 1629–1639. 10.1093/icb/icac033

34. Chapman, J. R., Dowell, M. A., Chan, R., & Unckless, R. L. (2020). The Genetic Basis of Natural Variation in *Drosophila melanogaster* Immune Defense against *Enterococcus faecalis*. Genes, 11(2), 234. 10.3390/genes11020234

35. Cociancich, S., Ghazi, A., Hetru, C., Hoffmann, J. A., & Letellier, L. (1993). Insect defensin, an inducible antibacterial peptide, forms voltage-dependent channels in *Micrococcus luteus*. Journal of Biological Chemistry, 268(26), 19239–19245. 10.1016/S0021-9258(19)36505-6

36. Cumnock, K., Gupta, A. S., Lissner, M., Chevee, V., Davis, N. M., & Schneider, D. S. (2018). Host Energy Source Is Important for Disease Tolerance to Malaria. Current Biology: CB, 28(10), 1635–1642.e3. 10.1016/j.cub.2018.04.009

37. De Gregorio, E., Spellman, P. T., Tzou, P., Rubin, G. M., & Lemaitre, B. (2002). The Toll and Imd pathways are the major regulators of the immune response in *Drosophila*. EMBO Journal, 21(11), 2568–2579. 10.1093/emboj/21.11.2568

38. DiAngelo, J. R., Bland, M. L., Bambina, S., Cherry, S., & Birnbaum, M. J. (2009). The immune response attenuates growth and nutrient storage in *Drosophila* by reducing insulin signaling. Proceedings of the National Academy of Sciences, 106(49), 20853–20858. 10.1073/pnas.0906749106

39. Diard, M., & Hardt, W.-D. (2017). Evolution of bacterial virulence. FEMS Microbiology Reviews, 41(5), 679–697. 10.1093/femsre/fux023

40. Dieppois, G., Opota, O., Lalucat, J., & Lemaitre, B. (2015). *Pseudomonas entomophila*: A Versatile Bacterium with Entomopathogenic Properties. In J.-L. Ramos, J. B. Goldberg, & A. Filloux (Eds.), Pseudomonas: Volume 7: New Aspects of Pseudomonas Biology (pp. 25–49). Springer Netherlands. 10.1007/978-94-017-9555-5_2

41. Dionne, M. (2014). Immune-metabolic interaction in *Drosophila*. Fly, 8(2), 75–79. 10.4161/fly.28113

42. Doeschl-Wilson, A. B., Bishop, S., Kyriazakis, I., & Villanueva, B. (2012). Novel methods for quantifying individual host response to infectious pathogens for genetic analyses. Frontiers in Genetics, 3. 10.3389/fgene.2012.00266

43. Duneau, D. F., Kondolf, H. C., Im, J. H., Ortiz, G. A., Chow, C., Fox, M. A., Eugénio, A. T., Revah, J., Buchon, N., & Lazzaro, B. P. (2017). The Toll pathway underlies host sexual dimorphism in resistance to both Gram-negative and Gram-positive bacteria in mated *Drosophila*. BMC Biology, 15(1), 124. 10.1186/s12915-017-0466-3

44. Duneau, D., Ferdy, J. B., Revah, J., Kondolf, H., Ortiz, G. A., Lazzaro, B. P., & Buchon, N. (2017). Stochastic variation in the initial phase of bacterial infection predicts the probability of survival in *D. melanogaster*. eLife, 6. 10.7554/eLife.28298

45. Duneau, D., & Ferdy, J.-B. (2022). Pathogen within-host dynamics and disease outcome: What can we learn from insect studies? Current Opinion in Insect Science, 100925. 10.1016/J.COIS.2022.100925

46. Duneau, D., Lafont, P. D. M., Lauzeral, C., Parthuisot, N., Faucher, C., Jin, X., Buchon, N., & Ferdy, J.-B. (2024). A within-host infection model to explore tolerance and resistance (p. 2021.10.19.464998). bioRxiv. 10.1101/2021.10.19.464998

47. Fabian, D. K., Garschall, K., Klepsatel, P., Santos-Matos, G., Sucena, É., Kapun, M., Lemaitre, B., Schlötterer, C., Arking, R., & Flatt, T. (2018). Evolution of longevity improves immunity in *Drosophila*. Evolution Letters, 2(6), 567–579. 10.1002/evl3.89

48. Faria, V. G., Martins, N. E., Paulo, T., Teixeira, L., Sucena, É., & Magalhães, S. (2015). Evolution of *Drosophila* resistance against different pathogens and infection routes entails no detectable maintenance costs. Evolution, 69(11), 2799–2809. 10.1111/evo.12782

49. Fehlbaum, P., Bulet, P., Michaut, L., Lagueux, M., Broekaert, W. F., Hetru, C., & Hoffmann, J. A. (1994). Insect immunity. Septic injury of *Drosophila* induces the synthesis of a potent antifungal peptide with sequence homology to plant antifungal peptides. Journal of Biological Chemistry, 269(52), 33159–33163. 10.1016/S0021-9258(20)30111-3

50. Ferrandon, D., Jung, A. C., Criqui, M., Lemaitre, B., Uttenweiler-Joseph, S., Michaut, L., Reichhart, J., & Hoffmann, J. A. (1998). A drosomycin-GFP reporter transgene reveals a local immune response in *Drosophila* that is not dependent on the Toll pathway. The EMBO Journal, 17(5), 1217–1227. 10.1093/emboj/17.5.1217

51. French, S. S., DeNardo, D. F., & Moore, M. C. (2007). Trade-offs between the reproductive and immune systems: Facultative responses to resources or obligate responses to reproduction? The American Naturalist, 170(1), 79–89. 10.1086/518569

52. Garschall, K., & Flatt, T. (2018). The interplay between immunity and aging in *Drosophila*. F1000Research, 7. 10.12688/f1000research.13117.1

53. Gates, D. E., Staley, M., Tardy, L., Giraudeau, M., Hill, G. E., McGraw, K. J., & Bonneaud, C. (2021). Levels of pathogen virulence and host resistance both shape the antibody response to an emerging bacterial disease. Scientific Reports, 11(1), 8209. 10.1038/s41598-021-87464-9

54. Gupta, V., Stewart, C. O., Rund, S. S. C., Monteith, K., & Vale, P. F. (2017). Costs and benefits of sublethal Drosophila C virus infection. Journal of Evolutionary Biology, 30(7), 1325–1335. 10.1111/jeb.13096

55. Gupta, V., Vasanthakrishnan, R. B., Siva-Jothy, J., Monteith, K. M., Brown, S. P., & Vale, P. F. (2017). The route of infection determines *Wolbachia* antibacterial protection in *Drosophila*. Proceedings of the Royal Society B: Biological Sciences, 284(1856), 20170809. 10.1098/rspb.2017.0809

56. Ha, E. M., Oh, C. T., Bae, Y. S., & Lee, W. J. (2005). A direct role for dual oxidase in *Drosophila* gut immunity. Science, 310(5749), 847–850. 10.1126/science.1117311

57. Hanson, M. A., Dostálová, A., Ceroni, C., Poidevin, M., Kondo, S., & Lemaitre, B. (2019). Synergy and remarkable specificity of antimicrobial peptides in vivo using a systematic knockout approach. eLife, 8, e44341. 10.7554/eLife.44341

58. Hanson, M. A., & Lemaitre, B. (2020). New insights on *Drosophila* antimicrobial peptide function in host defense and beyond. Current Opinion in Immunology, 62. 10.1016/j.coi.2019.11.008

59. Hart, B. L., & Hart, L. A. (2018). How mammals stay healthy in nature: The evolution of behaviours to avoid parasites and pathogens. Philosophical Transactions of the Royal Society B: Biological Sciences, 373(1751), 20170205. 10.1098/rstb.2017.0205

60. Hite, J. L., Pfenning-Butterworth, A., Stuart, ·, & Auld, K. J. R. (2023). Commentary: Infectious disease—The ecological theater and the evolutionary play. Evolutionary Ecology 2023, 1–11. 10.1007/S10682-023-10229-5

61. Hoffmann, J. A. (2003). The immune response of *Drosophila*. Nature 2003 426:6962, 426(6962), 33–38. 10.1038/nature02021

62. Hong, S., Sun, Y., Sun, D., & Wang, C. (2022). Microbiome assembly on *Drosophila* body surfaces benefits the flies to combat fungal infections. iScience, 25(6), 104408. 10.1016/j.isci.2022.104408

63. Horrocks, N. P. C., Matson, K. D., & Tieleman, B. I. (2011). Pathogen Pressure Puts Immune Defense into Perspective. Integrative and Comparative Biology, 51(4), 563–576. 10.1093/icb/icr011

64. Howick, V. M., & Lazzaro, B. P. (2017). The genetic architecture of defence as resistance to and tolerance of bacterial infection in *Drosophila melanogaster*. Molecular Ecology, 26(6), 1533–1546. 10.1111/mec.14017

65. Hudry, B., Khadayate, S., & Miguel-Aliaga, I. (2016). The sexual identity of adult intestinal stem cells controls organ size and plasticity. Nature, 530(7590), 344–348. 10.1038/nature16953

66. Hultmark, D. (2003). *Drosophila* immunity: Paths and patterns. Current Opinion in Immunology, 15(1), 12–19. 10.1016/s0952-7915(02)00005-5

67. Hultmark, D., Engström, A., Andersson, K., Steiner, H., Bennich, H., & Boman, H. G. (1983). Insect immunity. Attacins, a family of antibacterial proteins from *Hyalophora cecropia*. The EMBO Journal, 2(4), 571–576. 10.1002/j.1460-2075.1983.tb01465.x

68. Jiang, H., Patel, P. H., Kohlmaier, A., Grenley, M. O., McEwen, D. G., & Edgar, B. A. (2009). Cytokine/Jak/Stat signaling mediates regeneration and homeostasis in the *Drosophila* midgut. Cell, 137(7), 1343–1355. 10.1016/j.cell.2009.05.014

69. Jones, R. M., Luo, L., Ardita, C. S., Richardson, A. N., Kwon, Y. M., Mercante, J. W., Alam, A., Gates, C. L., Wu, H., Swanson, P. A., Lambeth, J. D., Denning, P. W., & Neish, A. S. (2013). Symbiotic lactobacilli stimulate gut epithelial proliferation via Nox-mediated generation of reactive oxygen species. The EMBO Journal, 32(23), 3017–3028. 10.1038/emboj.2013.224

70. Jung, Y.-J., Lee, Y.-T., Ngo, V. L., Cho, Y.-H., Ko, E.-J., Hong, S.-M., Kim, K.-H., Jang, J.-H., Oh, J.-S., Park, M.-K., Kim, C.-H., Sun, J., & Kang, S.-M. (2017). Heat-killed *Lactobacillus casei* confers broad protection against influenza A virus primary infection and develops heterosubtypic immunity against future secondary infection. Scientific Reports, 7(1), 17360. 10.1038/s41598-017-17487-8

71. Kawase, M., He, F., Kubota, A., Yoda, K., Miyazawa, K., & Hiramatsu, M. (2012). Heat-killed *Lactobacillus gasseri* TMC0356 protects mice against influenza virus infection by stimulating gut and respiratory immune responses. FEMS Immunology & Medical Microbiology, 64(2), 280–288. 10.1111/j.1574-695X.2011.00903.x

72. Keebaugh, E. S., Yamada, R., Obadia, B., Ludington, W. B., & Ja, W. W. (2018). Microbial Quantity Impacts *Drosophila* Nutrition, Development, and Lifespan. iScience, 4, 247–259. 10.1016/J.ISCI.2018.06.004

73. Kleino, A., Myllymäki, H., Kallio, J., Vanha-aho, L.-M., Oksanen, K., Ulvila, J., Hultmark, D., Valanne, S., & Rämet, M. (2008). *Pirk* Is a Negative Regulator of the *Drosophila* Imd Pathway. The Journal of Immunology, 180(8), 5413– 5422. 10.4049/JIMMUNOL.180.8.5413

74. Kotas, M. E., & Medzhitov, R. (2015). Homeostasis, Inflammation, and Disease Susceptibility. Cell, 160(5), 816–827. 10.1016/j.cell.2015.02.010

75. Kuraishi, T., Binggeli, O., Opota, O., Buchon, N., & Lemaitre, B. (2011). Genetic evidence for a protective role of the peritrophic matrix against intestinal bacterial infection in *Drosophila melanogaster*. Proceedings of the National Academy of Sciences of the United States of America, 108(38), 15966– 15971. 10.1073/pnas.1105994108

76. Kuraishi, T., Hori, A., & Kurata, S. (2013). Host-microbe interactions in the gut of *Drosophila melanogaster*. Frontiers in Physiology, 4. 10.3389/fphys.2013.00375

77. Kutzer, M. A. M., & Armitage, S. A. O. (2016). Maximising fitness in the face of parasites: A review of host tolerance. Zoology, 119(4), 281–289. 10.1016/j.zool.2016.05.011

78. Kutzer, M. A. M., Gupta, V., Neophytou, K., Doublet, V., Monteith, K. M., & Vale, P. F. (2023). Intraspecific genetic variation in host vigour, viral load and disease tolerance during Drosophila C virus infection. Open Biology, 13(3), 230025. 10.1098/RSOB.230025

79. Kutzer, M. A. M., Kurtz, J., & Armitage, S. A. O. (2018). Genotype and diet affect resistance, survival, and fecundity but not fecundity tolerance. Journal of Evolutionary Biology, 31(1), 159–171. 10.1111/jeb.13211

80. Kuznetsova, A., Brockhoff, P. B., & Christensen, R. H. B. (2017). lmerTest Package: Tests in Linear Mixed Effects Models. Journal of Statistical Software, 82, 1–26. 10.18637/jss.v082.i13

81. Lambert, J., DUNBARt, B., Lepage, P., Dorsselaer, A. V., Hoffmann, J., Fothergilli, J., & Hoffmann, Dani. (1989). Insect immunity: Isolation from immune blood of the dipteran *Phormia terranovae* of two insect antibacterial peptides with sequence homology to rabbit lung macrophage bactericidal peptides. Proc. Natl. Acad. Sci. USA.

82. Lazzaro, B. P., & Tate, A. T. (2022). Balancing sensitivity, risk, and immunopathology in immune regulation. Current Opinion in Insect Science, 100874. 10.1016/J.COIS.2022.100874

83. Le Bourg, É. (2011). A cold stress applied at various ages can increase resistance to heat and fungal infection in aged *Drosophila melanogaster* flies. Biogerontology, 12(3), 185–193. 10.1007/s10522-010-9309-0

84. Lee, K.-A., Kim, S.-H., Kim, E.-K., Ha, E.-M., You, H., Kim, B., Kim, M.-J., Kwon, Y., Ryu, J.-H., & Lee, W.-J. (2013). Bacterial-derived uracil as a modulator of mucosal immunity and gut-microbe homeostasis in *Drosophila*. Cell, 153(4), 797–811. 10.1016/j.cell.2013.04.009

85. Lemaitre, B. (2015). *Pseudomonas entomophila*: A versatile bacterium with entomopathogenic properties. Pseudomonas: Volume 7: New Aspects of Pseudomonas Biology, 25–49. 10.1007/978-94-017-9555-5_2

86. Lemaitre, B., & Hoffmann, J. (2007). The Host Defense of *Drosophila melanogaster*. Annual Review of Immunology, 25(1), 697–743. 10.1146/annurev.immunol.25.022106.141615

87. Lemaitre, B., & Miguel-Aliaga, I. (2013). The Digestive Tract of *Drosophila melanogaster*. Annu Rev Genet., 47, 377–404. 10.1146/annurev-genet-111212-133343

88. Lemaitre, B., Reichhart, J.-M., & Hoffmann, J. A. (1997). *Drosophila* host defense: Differential induction of antimicrobial peptide genes after infection by various classes of microorganisms. Proceedings of the National Academy of Sciences, 94(26), 14614–14619. 10.1073/pnas.94.26.14614

89. Lenth, R. V. (2017). emmeans: Estimated Marginal Means, aka Least-Squares Means (p. 1.10.3). 10.32614/CRAN.package.emmeans

90. Li, X., Rommelaere, S., Kondo, S., & Lemaitre, B. (2020). Renal Purge of Hemolymphatic Lipids Prevents the Accumulation of ROS-Induced Inflammatory Oxidized Lipids and Protects *Drosophila* from Tissue Damage. Immunity, 52(2), 374–387.e6. 10.1016/J.IMMUNI.2020.01.008

91. Liehl, P., Blight, M., Vodovar, N., Boccard, F., & Lemaitre, B. (2006). Prevalence of local immune response against oral infection in a *Drosophila/Pseudomonas* infection model. PLoS Pathogens, 2(6), 0551–0561. 10.1371/journal.ppat.0020056

92. Lissner, M. M., & Schneider, D. S. (2018). The physiological basis of disease tolerance in insects. Current Opinion in Insect Science, 29, 133–136. 10.1016/j.cois.2018.09.004

93. Lopes, P. C., French, S. S., Woodhams, D. C., & Binning, S. A. (2022). Infection avoidance behaviors across vertebrate taxa: Patterns, processes, and future directions. In V. Ezenwa, S. M. Altizer, & R. Hall (Eds.), Animal Behavior and Parasitism (p. 0). Oxford University Press. 10.1093/oso/9780192895561.003.0014

94. Loudhaief, R., Jneid, R., Christensen, C. F., Mackay, D. J., Andersen, D. S., & Colombani, J. (2023). The *Drosophila* tumor necrosis factor receptor, *Wengen*, couples energy expenditure with gut immunity. Science Advances, 9(23), eadd4977. 10.1126/sciadv.add4977

95. Luckinbill, L. S., Arking, R., Clare, M. J., Cirocco, W. C., & Buck, S. A. (1984). Selection for delayed senescence in *Drosophila melanogaster*. Evolution; International Journal of Organic Evolution, 38(5), 996–1003. 10.1111/j.1558-5646.1984.tb00369.x

96. Martins, N. E., Faria, V. G., Teixeira, L., Magalhães, S., & Sucena, É. (2013). Host Adaptation Is Contingent upon the Infection Route Taken by Pathogens. PLoS Pathogens, 9(9), e1003601. 10.1371/journal.ppat.1003601

97. Martins, N. E., Faria, V. G., Nolte, V., Schlötterer, C., Teixeira, L., Sucena, É., & Magalhães, S. (2014). Host adaptation to viruses relies on few genes with different cross-resistance properties. Proceedings of the National Academy of Sciences of the United States of America, 111(16), 5938–5943. 10.1073/pnas.1400378111

98. Martins, R., Carlos, A. R., Braza, F., Thompson, J. A., Bastos-Amador, P., Ramos, S., & Soares, M. P. (2019). Disease Tolerance as an Inherent Component of Immunity. Annual Review of Immunology 37:405-437. 10.1146/annurev-immunol-042718

99. Mccarville, J. L., & Ayres, J. S. (2017). Disease tolerance: Concept and mechanisms. Curr Opin Immunol. 50:88–93. 10.1016/j.coi.2017.12.003

100. Medzhitov, R., Schneider, D. S., & Soares, M. P. (2012). Disease tolerance as a defense strategy. Science, 335(6071), 936–941. 10.1126/science.1214935

101. Meiselman, M. R., Kingan, T. G., & Adams, M. E. (2018). Stress-induced reproductive arrest in *Drosophila* occurs through ETH deficiency-mediated suppression of oogenesis and ovulation. BMC Biology, 16(1), 18. 10.1186/s12915-018-0484-9

102. Miguel-Aliaga, I., Jasper, H., & Lemaitre, B. (2018). Anatomy and Physiology of the Digestive Tract of *Drosophila melanogaster*. Genetics, 210(2):357–396 10.1534/genetics.118.300224

103. Minute, L., Bergón-Gutiérrez, M., Mata-Martínez, P., Fernández-Pascual, J., Terrón, V., Bravo-Robles, L., Bıçakcıoğlu, G., Zapata-Fernández, G., Aguiló, N., López-Collazo, E., & Del Fresno, C. (2024). Heat-killed *Mycobacterium tuberculosis* induces trained immunity in vitro and in vivo administered systemically or intranasally. iScience, 27(2), 108869. 10.1016/j.isci.2024.108869

104. Miyashita, A., Kizaki, H., Kawasaki, K., Sekimizu, K., & Kaito, C. (2014). Primed Immune Responses to Gram-negative Peptidoglycans Confer Infection Resistance in Silkworms. Journal of Biological Chemistry, 289(20), 14412– 14421. 10.1074/jbc.M113.525139

105. Miyashita, A., Takahashi, S., Ishii, K., Sekimizu, K., & Kaito, C. (2015). Primed Immune Responses Triggered by Ingested Bacteria Lead to Systemic Infection Tolerance in Silkworms. PLOS ONE, 10(6), e0130486. 10.1371/journal.pone.0130486

106. Myllymäki, H., & Rämet, M. (2014). JAK/STAT Pathway in Drosophila Immunity. Scandinavian Journal of Immunology, 79(6), 377–385. 10.1111/sji.12170

107. Nayak, N., & Mishra, M. (2022). *Drosophila melanogaster* as a model to understand the mechanisms of infection mediated neuroinflammation in neurodegenerative diseases. Journal of Integrative Neuroscience, 21(2), Article 2. 10.31083/j.jin2102066

108. Neyen, C., Bretscher, A. J., Binggeli, O., & Lemaitre, B. (2014). Methods to study *Drosophila* immunity. Methods, 68(1), 116–128. 10.1016/j.ymeth.2014.02.023

109. Neyen, C., Poidevin, M., Roussel, A., & Lemaitre, B. (2012). Tissue- and Ligand-Specific Sensing of Gram-Negative Infection in Drosophila by PGRP-LC Isoforms and PGRP-LE. The Journal of Immunology, 189(4), 1886–1897. 10.4049/jimmunol.1201022

110. Nixon, D. F., Kyza-Karavioti, M., Mallick, S., Daley, L., Hupert, N., Bachtel, N. D., & Eleftherianos, I. (2024). A model of proximate protection against pathogenic infection through shared immunity. mBio, 15(12), e03046–24. 10.1128/mbio.03046-24

111. Nonaka, S., Salim, E., Kamiya, K., Hori, A., Nainu, F., Asri, R. M., Masyita, A., Nishiuchi, T., Takeuchi, S., Kodera, N., & Kuraishi, T. (2020). Molecular and Functional Analysis of Pore-Forming Toxin Monalysin From Entomopathogenic Bacterium *Pseudomonas entomophila*. Frontiers in Immunology, 11, 520. 10.3389/fimmu.2020.00520

112. Nunes, C., Koyama, T., & Sucena, É. (2021). Co-option of immune effectors by the hormonal signalling system triggering metamorphosis in *Drosophila melanogaster*. PLOS Genetics, 17(11), e1009916. 10.1371/journal.pgen.1009916

113. Ohlstein, B., & Spradling, A. (2006). The adult *Drosophila* posterior midgut is maintained by pluripotent stem cells. Nature, 439(7075), 470–474. 10.1038/nature04333

114. Opota, O., Vallet-Gély, I., Vincentelli, R., Kellenberger, C., Iacovache, I., Gonzalez, M. R., Roussel, A., van der Goot, F. G., & Lemaitre, B. (2011). Monalysin, a novel ß-pore-forming toxin from the *Drosophila* pathogen *Pseudomonas entomophila*, contributes to host intestinal damage and lethality. PLoS Pathogens, 7(9). 10.1371/journal.ppat.1002259

115. Palmer, W. H., Medd, N. C., Beard, P. M., & Obbard, D. J. (2018). Isolation of a natural DNA virus of *Drosophila melanogaster*, and characterisation of host resistance and immune responses. PLoS Pathogens, 14(6). 10.1371/journal.ppat.1007050

116. Parker, D. J., Envall, T., Ritchie, M. G., & Kankare, M. (2020). Sex-specific responses to cold in a very cold-tolerant, northern *Drosophila* species. Heredity. 10.1038/s41437-020-00398-2

117. Paulo, T. F., Akyaw, P. A., Paixão, T., & Sucena, É. (2023). Adaptation to oral infection in D. melanogaster through evolution of both resistance and disease tolerance mechanisms (p. 2023.08.23.554397). bioRxiv. 10.1101/2023.08.23.554397

118. Prakash, A., Monteith, K. M., Bonnet, M., & Vale, P. F. (2023). Duox and Jak/Stat signalling influence disease tolerance in *Drosophila* during *Pseudomonas entomophila* infection. Developmental & Comparative Immunology, 147, 104756. 10.1016/J.DCI.2023.104756

119. Prakash, A., Monteith, K. M., & Vale, P. F. (2022). Mechanisms of damage prevention, signalling and repair impact disease tolerance. Proceedings of the Royal Society B: Biological Sciences, 289(1981). 10.1098/RSPB.2022.0837

120. Prakash, A., Monteith, K. M., & Vale, P. F. (2024). Negative immune regulation contributes to disease tolerance in *Drosophila melanogaster*. Physiological Entomology, n/a(n/a). 10.1111/phen.12464

121. Pursall, E. R., & Rolff, J. (2011). Immune Responses Accelerate Ageing: Proof-of-Principle in an Insect Model. PLOS ONE, 6(5), e19972. 10.1371/journal.pone.0019972

122. R Core Team. (2021). R: The R Project for Statistical Computing. https://www.r-project.org/

123. Råberg, L., Graham, A. L., & Read, A. F. (2009). Decomposing health: Tolerance and resistance to parasites in animals. Philos Trans R Soc Lond B Biol Sci. 364(1513):37–49. 10.1098/rstb.2008.0184

124. Råberg, L. (2014). How to live with the enemy: Understanding tolerance to parasites. PLoS Biology, 12(11), e1001989. 10.1371/journal.pbio.1001989

125. Råberg, L., Sim, D., & Read, A. F. (2007). Disentangling genetic variation for resistance and tolerance to infectious diseases in animals. Science, 318(5851), 812–814. 10.1126/science.1148526

126. Raval, D., Daley, L., & Eleftherianos, I. (2023). *Drosophila melanogaster* larvae are tolerant to oral infection with the bacterial pathogen *Photorhabdus luminescens*. microPublication Biology, 2023, 10.17912/micropub.biology.000938. https://doi.org/10.17912/micropub.biology.000938

127. Regan, J. C., Khericha, M., Dobson, A. J., Bolukbasi, E., Rattanavirotkul, N., & Partridge, L. (2016). Sex difference in pathology of the ageing gut mediates the greater response of female lifespan to dietary restriction. eLife, 5(February2016). 10.7554/eLife.10956

128. Regan, J. C., Lu, Y.-X., Ureña, E., Meilenbrock, R. L., Catterson, J. H., Kißler, D., Fröhlich, J., Funk, E., & Partridge, L. (2022). Sexual identity of enterocytes regulates autophagy to determine intestinal health, lifespan and responses to rapamycin. Nature Aging, 2(12), 1145–1158. 10.1038/s43587-022-00308-7

129. Regoes, R. R., McLaren, P. J., Battegay, M., Bernasconi, E., Calmy, A., Günthard, H. F., Hoffmann, M., Rauch, A., Telenti, A., Fellay, J., & Study, the S. H. C. (2014). Disentangling Human Tolerance and Resistance Against HIV. PLOS Biology, 12(9), e1001951. 10.1371/journal.pbio.1001951

130. Reiff, T., Jacobson, J., Cognigni, P., Antonello, Z., Ballesta, E., Tan, K. J., Yew, J. Y., Dominguez, M., & Miguel-Aliaga, I. (2015). Endocrine remodelling of the adult intestine sustains reproduction in *Drosophila*. eLife, 4, e06930. 10.7554/eLife.06930

131. Round, J. L., & Mazmanian, S. K. (2009). The gut microbiota shapes intestinal immune responses during health and disease. Nature Reviews Immunology, 9(5), 313–323. 10.1038/nri2515

132. Roy, B. A., & Kirchner, J. W. (2000). Evolutionary dynamics of pathogen resistance and tolerance. Evolution, 54(1), 51–63. 10.1111/j.0014-3820.2000.tb00007.x

133. Sadd, B. M., & Schmid-Hempel, P. (2009). PERSPECTIVE: Principles of ecological immunology. Evolutionary Applications, 2(1), 113–121. 10.1111/j.1752-4571.2008.00057.x

134. Sadd, B. M., & Siva-Jothy, M. T. (2006). Self-harm caused by an insect’s innate immunity. Proceedings of the Royal Society B: Biological Sciences, 273(1600), 2571–2574. 10.1098/rspb.2006.3574

135. Sarabian, C., Curtis, V., & McMullan, R. (2018). Evolution of pathogen and parasite avoidance behaviours. *Philosophical Transactions of the Royal Society of London. Series B*, Biological Sciences, 373(1751), 20170256. 10.1098/rstb.2017.0256

136. Sartor, R. B. (2008). Microbial influences in inflammatory bowel diseases. Gastroenterology, 134(2), 577–594. 10.1053/j.gastro.2007.11.059

137. Schlamp, F., Delbare, S. Y. N., Early, A. M., Wells, M. T., Basu, S., & Clark, A. G. (2021). Dense time-course gene expression profiling of the *Drosophila melanogaster* innate immune response. BMC Genomics 2021 22:1, 22(1), 1–22. 10.1186/S12864-021-07593-3

138. Schmid-Hempel, P. (2003). Variation in immune defence as a question of evolutionary ecology. Proceedings of the Royal Society of London. Series B: Biological Sciences, 270(1513), 357–366. 10.1098/rspb.2002.2265

139. Schneider, D. S., & Ayres, J. S. (2008). Two ways to survive infection: What resistance and tolerance can teach us about treating infectious diseases. Nature Reviews Immunology, 8(11), 889–895. 10.1038/nri2432

140. Schulenburg, H., Kurtz, J., Moret, Y., & Siva-Jothy, M. T. (2008). Introduction. Ecological immunology. Philosophical Transactions of the Royal Society B: Biological Sciences, 364(1513), 3–14. 10.1098/rstb.2008.0249

141. Schwenke, R. A., Lazzaro, B. P., & Wolfner, M. F. (2016). Reproduction–Immunity Trade-Offs in Insects. Annual Review of Entomology, 61(1), 239–256. 10.1146/annurev-ento-010715-023924

142. Shahrestani, P., Chambers, M., Vandenberg, J., Garcia, K., Malaret, G., Chowdhury, P., Estrella, Y., Zhu, M., & Lazzaro, B. P. (2018). Sexual dimorphism in *Drosophila melanogaster* survival of *Beauveria bassiana* infection depends on core immune signaling. Scientific Reports, 8(1), 12501. 10.1038/s41598-018-30527-1

143. Simms, E. L. (2000). Defining tolerance as a norm of reaction. Evolutionary Ecology, 14(4), 563–570. 10.1023/A:1010956716539

144. Soares, M. P., Gozzelino, R., & Weis, S. (2014). Tissue damage control in disease tolerance. Trends in Immunology, 35(10), 483–494. 10.1016/j.it.2014.08.001

145. Socha, C., Pais, I. S., Lee, K.-Z., Liu, J., Liégeois, S., Lestradet, M., & Ferrandon, D. (2023). Fast *Drosophila* enterocyte regrowth after infection involves a reverse metabolic flux driven by an amino acid transporter. iScience, 26(9), 107490. 10.1016/j.isci.2023.107490

146. Stearns, S. C., & Stearns, S. C. (1992). The Evolution of Life Histories. Oxford University Press.

147. Sun, S.-C., & Faye, I. (1992). Cecropia immunoresponsive factor, an insect immunoresponsive factor with DNA-binding properties similar to nuclear-factor KB. European Journal of Biochemistry, 204(2), 885–892. 10.1111/j.1432-1033.1992.tb16708.x

148. Syed, Z. A., Gupta, V., Arun, M. G., Dhiman, A., Nandy, B., & Prasad, N. G. (2020). Absence of reproduction-immunity trade-off in male *Drosophila melanogaster* evolving under differential sexual selection. BMC Evolutionary Biology, 20(1), 13. 10.1186/s12862-019-1574-1

149. Tafesh-Edwards, G., & Eleftherianos, I. (2023). The role of *Drosophila* microbiota in gut homeostasis and immunity. Gut Microbes, 15(1), 2208503. 10.1080/19490976.2023.2208503

150. Teder, T., & Tammaru, T. (2005). Sexual size dimorphism within species increases with body size in insects. Oikos, 108(2), 321–334. 10.1111/j.0030-1299.2005.13609.x

151. Troha, K., & Buchon, N. (2019). Methods for the study of innate immunity in *Drosophila melanogaster*. Wiley Interdisciplinary Reviews: Developmental Biology, 8(5), 1–25. 10.1002/wdev.344

152. Troha, K., Im, J. H., Revah, J., Lazzaro, B. P., & Buchon, N. (2018). Comparative transcriptomics reveals *CrebA* as a novel regulator of infection tolerance in *D. melanogaster*. PLOS Pathogens, 14(2), e1006847. 10.1371/journal.ppat.1006847

153. Tzou, P., Ohresser, S., Ferrandon, D., Capovilla, M., Reichhart, J. M., Lemaitre, B., Hoffmann, J. A., & Imler, J. L. (2000). Tissue-specific inducible expression of antimicrobial peptide genes in *Drosophila* surface epithelia. Immunity, 13(5), 737–748. 10.1016/s1074-7613(00)00072-8

154. Valanne, S., Wang, J.-H., & Rämet, M. (2011). The *Drosophila* Toll Signaling Pathway. The Journal of Immunology, 186(2), 649–656. 10.4049/jimmunol.1002302

155. Vallet-Gely, I., Opota, O., Boniface, A., Novikov, A., & Lemaitre, B. (2010). A secondary metabolite acting as a signalling molecule controls *Pseudomonas entomophila* virulence. Cellular Microbiology, 12(11), 1666–1679. 10.1111/j.1462-5822.2010.01501.x

156. Villafranca, N., Changsut, I., Diaz de Villegas, S., Womack, H., & Fuess, L. E. (2023). Characterization of trade-offs between immunity and reproduction in the coral species *Astrangia poculata*. PeerJ, 11, e16586. 10.7717/peerj.16586

157. Vincent, C. M., Beckwith, E. J., Silva, C. J. S. da, Pearson, W. H., Kierdorf, K., Gilestro, G. F., & Dionne, M. S. (2022). Infection increases activity via Toll dependent and independent mechanisms in *Drosophila melanogaster*. PLOS Pathogens, 18(9), e1010826. 10.1371/journal.ppat.1010826

158. Vincent, C. M., & Dionne, M. S. (2021). Disparate regulation of IMD signaling drives sex differences in infection pathology in *Drosophila melanogaster*. Proceedings of the National Academy of Sciences, 118(32), e2026554118. 10.1073/PNAS.2026554118

159. Vodovar, N., Vallenet, D., Cruveiller, S., Rouy, Z., Barbe, V., Acosta, C., Cattolico, L., Jubin, C., Lajus, A., Segurens, B., Benoı^t Vacherie, B., Wincker, P., Weissenbach, J., Lemaitre, B., Médigue, C., & Boccard, F. (2006). Complete genome sequence of the entomopathogenic and metabolically versatile soil bacterium *Pseudomonas entomophila*. Nature Biotechnology 24(6), 3. 10.1038/nbt1212

160. Vodovar, N., Vinals, M., Liehl, P., Basset, A., Degrouard, J., Spellman, P., Boccard, F., & Lemaitre, B. (2005). *Drosophila* host defense after oral infection by an entomopathogenic *Pseudomonas* species. Proceedings of the National Academy of Sciences of the United States of America, 102(32), 11414– 11419. 10.1073/pnas.0502240102

161. Wen, Y., He, Z., Xu, T., Jiao, Y., Liu, X., Wang, Y. F., & Yu, X. Q. (2019). Ingestion of killed bacteria activates antimicrobial peptide genes in *Drosophila melanogaster* and protects flies from septic infection. Developmental and Comparative Immunology, 95, 10–18. 10.1016/j.dci.2019.02.001

162. Westlake, H., Hanson, A. M., & Lemaitre, B. (2024). The Drosophila immunity handbook (1st ed.). 10.55430/6304TDIHVA01

163. Whitlock, G. C., Lukaszewski, R. A., Judy, B. M., Paessler, S., Torres, A. G., & Estes, D. M. (2008). Host immunity in the protective response to vaccination with heat-killed *Burkholderia mallei*. BMC Immunology, 9(1), 55. 10.1186/1471-2172-9-55

164. Wickham, H. (2016). Ggplot2. Springer International Publishing. 10.1007/978-3-319-24277-4

165. Wickham, H., Averick, M., Bryan, J., Chang, W., McGowan, L. D., François, R., Grolemund, G., Hayes, A., Henry, L., Hester, J., Kuhn, M., Pedersen, T. L., Miller, E., Bache, S. M., Müller, K., Ooms, J., Robinson, D., Seidel, D. P., Spinu, V., … Yutani, H. (2019). Welcome to the Tidyverse. Journal of Open Source Software, 4(43), 1686. 10.21105/joss.01686

166. Zeileis, A., Kleiber, C., & Jackman, S. (2008). Regression Models for Count Data in R. Journal of Statistical Software, 27, 1–25. 10.18637/jss.v027.i08

167. Zélé, F., Santos-Matos, G., Figueiredo, A. R. T., Eira, C., Pinto, C., Laurentino, T. G., Sucena, É., & Magalhães, S. (2019). Spider mites escape bacterial infection by avoiding contaminated food. Oecologia, 189(1), 111–122. 10.1007/s00442-018-4316-y

168. Zeng, T., Jaffar, S., Xu, Y., & Qi, Y. (2022). The Intestinal Immune Defense System in Insects. Int. J. Mol. Sci, 2022, 15132. 10.3390/ijms232315132

169. Zerofsky, M., Harel, E., Silverman, N., & Tatar, M. (2005). Aging of the innate immune response in *Drosophila melanogaster*. Aging Cell, 4(2), 103–108. 10.1111/j.1474-9728.2005.00147.x

170. Zhai, Z., Huang, X., & Yin, Y. (2018). Beyond immunity: The Imd pathway as a coordinator of host defense, organismal physiology and behavior. Developmental and Comparative Immunology, 83, 51–59. 10.1016/j.dci.2017.11.008

171. Zhou, F., Rasmussen, A., Lee, S., & Agaisse, H. (2013). The Upd3 cytokine couples environmental challenge and intestinal stem cell division through modulation of JAK/STAT signaling in the stem cell microenvironment. Developmental Biology, 373(2), 383–393. 10.1016/j.ydbio.2012.10.023

172. Zhou, S. O., Arunkumar, R., Irfan, A., Ding, S. D., Leitão, A. B., & Jiggins, F. M. (2024). The evolution of constitutively active humoral immune defenses in *Drosophila* populations under high parasite pressure. PLOS Pathogens, 20(1), e1011729. 10.1371/journal.ppat.1011729

