## Supplementary Figures for "Pathogen-Induced Damage in Drosophila: Uncoupling Disease Tolerance from Resistance"

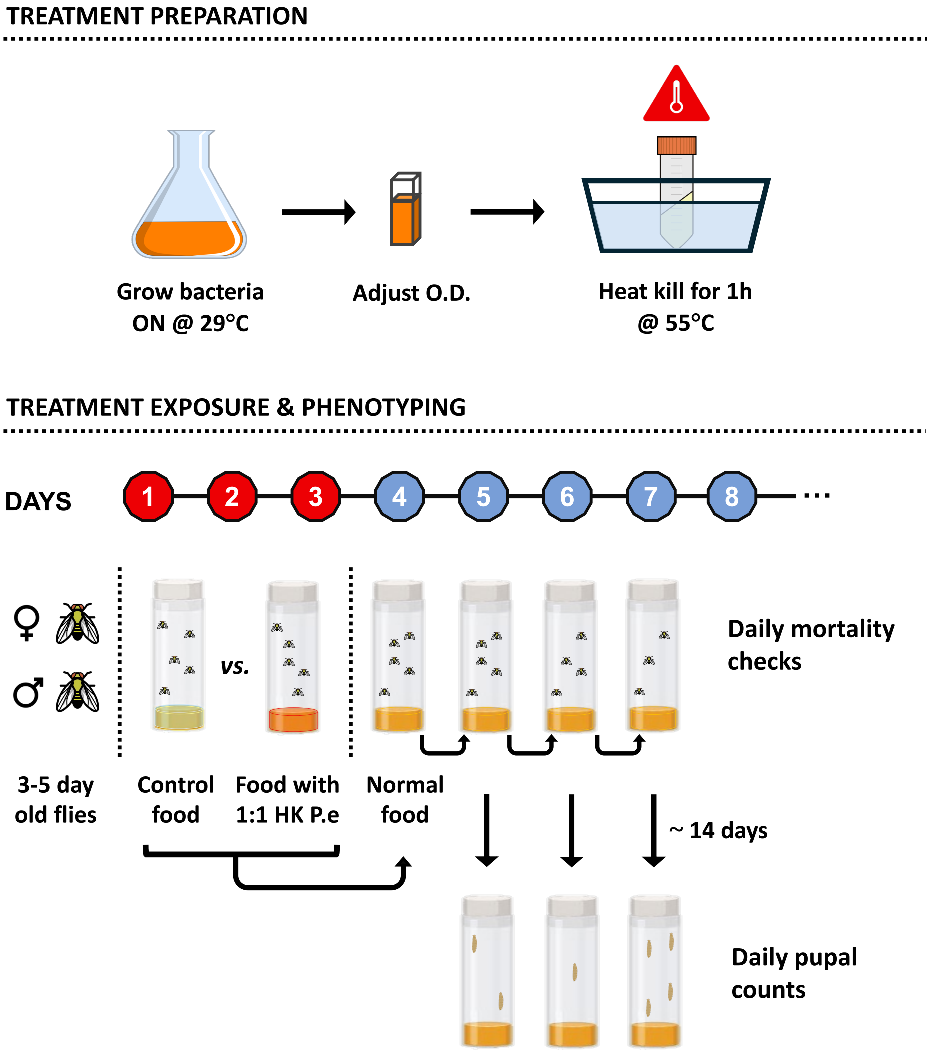


**Figure S1. Experimental setup for measuring survival and reproductive output of flies using Heat-killed *P. entomophila*.** Bacteria were grown under standard conditions (29°C at 180 RPM) and the OD was adjusted accordingly (OD600= 100), followed by a water bath incubation at 55°C for one hour, and freezing until later usage. Three to five day-old flies were either exposed to fly food mixed 1:1 with HK *P. entomophila* or with PBS (control food) for three days and monitored daily for survival. After three days, flies were placed on normal food and flipped daily into new vials for 12 to 18 days. Survival and pupal counts were measured in each of the vials where flies were maintained. Red circles indicate the period of exposure to food mixed with HK *P. entomophila*, while blue circles indicate periods where flies were kept on normal food.

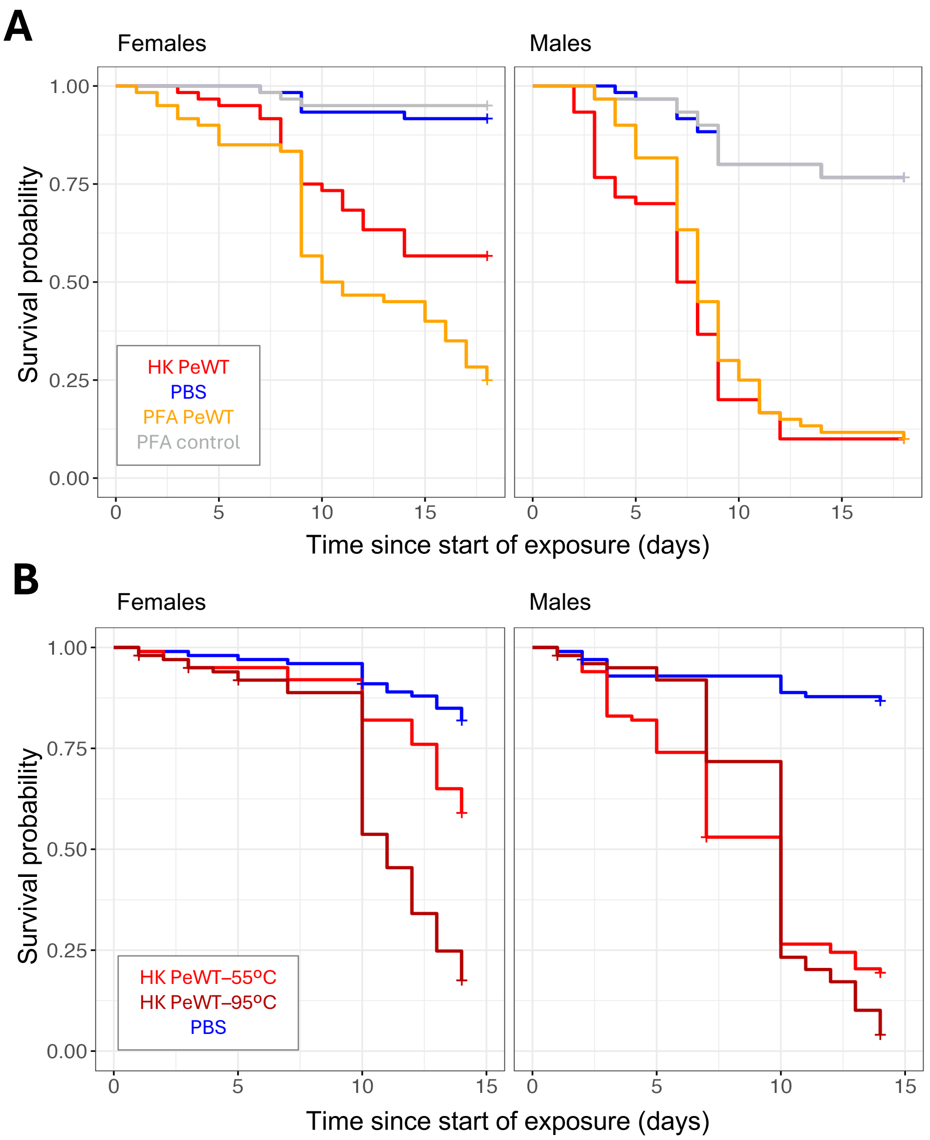


**Figure S2. Alternative inactivation methods confirm effect on survival.** Survival over 14 days of female (left plots) and male (right plots) flies exposed to food containing wild-type *P. entomophila* inactivated with different methods. **A)** Survival upon exposure to food containing *P. entomophila* heat-killed at 55ºC (HK PeWT - red) or fixed with paraformaldehyde (PFA) (PFA PeWT - orange), containing PFA alone (PFA control - grey) or PBS (PBS control - blue). There is no significant difference between survival measurements in the group fed with HK *P. entomophila* and that of PFA control ($p>0.05$), but both are different from control treatments ($p<0.001$). **B)** Survival upon exposure to food with heat-killed (HK) *P. entomophila* at 55ºC (HK PeWT-55ºC - red) or 95ºC (HK PeWT-95ºC - brown) or PBS control food (PBS - blue). There were differences in survival between inactivation temperatures in females ($p<0.001$), but not in males ($p>0.05$). Full details and outputs of statistical analyses can be found in Tables S1 and S2.

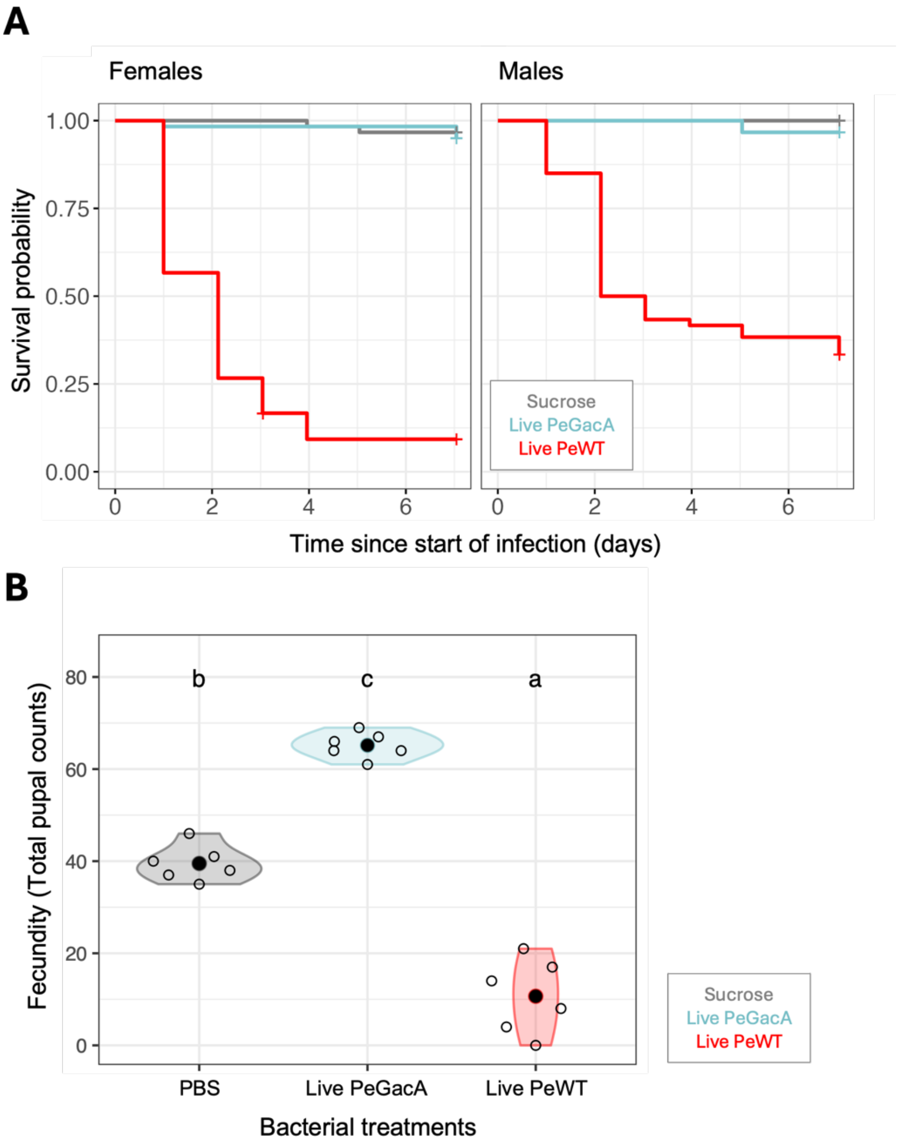


**Figure S3. Oral infection with wildtype *P. entomophila* leads to increased mortality and reduced fecundity in *D. melanogaster*.** Survival and fecundity in adult flies orally infected with either wild-type *P. entomophila* (PeWT - red), the avirulent *P. entomophila* $\Delta$*GacA* mutant (PeGacA - light blue), or 5% sucrose control solution (Sucrose - grey). **A)** Survival curves of females (left panel) and males (right panel) over 6 days show a significant difference in survival between the PeWT- and the GacA-infected or the sucrose individuals($p<0.001$ in both cases). **B)** Fecunidty ( measured as cumulative daily pupal counts; see Material and Methods) shows that flies infected with wildtype PeWT are significantly less fecund than the PeGacA-infected and the sucrose group ($p<0.001$ in both cases). PeGacA-infected flies also showed a significantly higher reproductive output than flies from the Sucrose group (STAT).Differences between groups were estimated by post hoc comparisons (Tukey's honest significant differences) and are indicated by different letters in each plot ($p<0.05$). Full details and outputs of statistical analyses can be found in Tables S1 and S6.

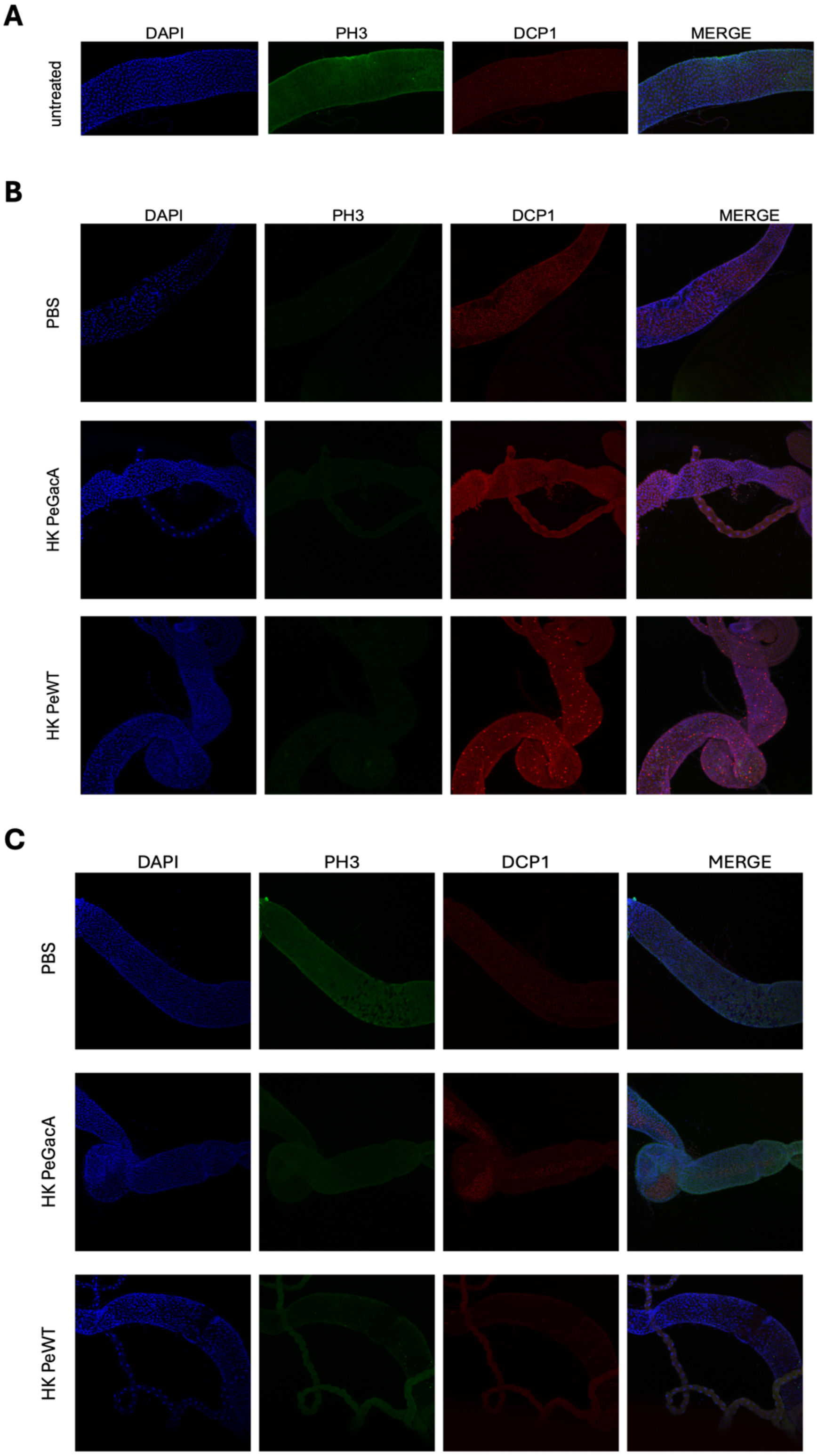


**Figure S4. Feeding on heat-killed *P. entomophila* induces apoptosis and mitotic cell division in gut epithelial tissues.** Immunofluorescent staining using antibodies against Drosophila caspase 1 (Dcp-1) in red, Phospho-H3 (PH3) in green and merged with DAPI in blue. Images are taken from the posterior midgut of males untreated (**A**) 72h **(B)** and 120h **(C),** after treatment with heat-killed wildtype (HK PeWT) or mutant ∆GacA (HK PeGacA) *P. entomophila*. Images are representatives of at least six replicates per treatment and time point, displayed according to channels and merged in the final column.
